# Combined Lytic Score as a new metric to quantify phage-host specificity, tested in a screening study on a collection of *Salmonella* strains

**DOI:** 10.64898/2026.01.28.702309

**Authors:** Michał Różański, Natalia Adamiak, Karolina Pospiech, Katarzyna Grochala, Jolanta Witaszewska, Justyna Matczak, Ewelina Wójcik, Jarosław Dastych

## Abstract

Bacteriophages are promising alternatives for antibiotics. One challenge of developing bacteriophage-based preparation for practical application is to understand expected host range of such products. We evaluated the host range and lytic activity of BAFASAL®, a four-phage cocktail targeting Salmonella in poultry production. For that purpose, we developed a new composite metric, the Combined Lytic Score (CLS), integrating results from two in vitro assays: serial dilutions spot test on semisolid medium and spectrophotometric growth inhibition in liquid culture. Using this approach phage cocktail was tested against collection of 72 Salmonella strains, including 55 S. Enteritidis isolates representing diverse geographic origins and genomic backgrounds. Spot test patterns were transformed into a continuous scale using the Most Probable Number (MPN) approach to estimate the number of phages required for visible lysis. In parallel, growth inhibition was quantified as the area-under-curve–based inhibition score (ANS). Both metrics were normalized and combined into CLS as a projection onto the regression line describing their correlation (R² ≈ 0.82). More than 65% of S. Enteritidis strains, reached normalized CLS values higher 75%, indicating high susceptibility to BAFASAL® in vitro. Phage susceptibility did not correlate with either phenotypic antibiotic resistance or the number of resistance and virulence genes. CLS provides a quantitative method to integrate different experimental methods of determination of bacterial susceptibility to bacteriophages and to rank bacterial strains by phage susceptibility. This approach supports robust host range determination and may facilitate regulatory evaluation and rational design of phage-based interventions in food safety and animal production.

**IMPORTANCE:** Assessment of bacteriophage host range is an important step in characterization of bacteriophage strains both in basic and translational research, yet it is still commonly based on qualitative or poorly standardized assays. This lack of harmonization limits reproducibility and complicates comparisons across studies, laboratories, and application contexts. In this work, we propose the Combined Lytic Score (CLS) as a quantitative framework that integrates outcomes from two widely used experimental approaches: serial-dilution spot assays and microtiter-based growth inhibition kinetics. By converting spot-test results into a continuous, concentration-dependent metric and combining them with normalized kinetic inhibition data, CLS enables more consistent interpretation of phage–host interaction outcomes. Application of CLS to a diverse collection of Salmonella enterica strains demonstrates how this approach can support systematic, scalable host range analyses. The CLS framework provides a practical step toward improved standardization of phage susceptibility testing, facilitating clearer data interpretation and comparison in both environmental and applied microbiology research.

## INTRODUCTION

Bacteriophages, or phages, are viruses that specifically infect and lyse bacterial cells. Due to their high specificity and bactericidal potential, phages are increasingly being explored as viable alternatives to antibiotics, particularly addressing the growing crisis of antimicrobial resistance (AMR). One of the most critical parameters defining a phage’s utility in clinical, veterinary, agricultural, or industrial applications is its host range—the spectrum of bacterial strains, species, or even genera that the phage is able to effectively infect and kill [1, 2, 3, 4, 5].

Host range is not a rigid category but rather a continuum, and although it is commonly defined as the phage’s ability to replicate and kill the host, Hyman et al. [6] proposed a set of terms with precise definitions based on the depth of phage-host interaction. For example, *adsorption range* refers to the ability of the phage to bind specifically to a receptor on the bacterial surface. *Penetration range* denotes the capacity to inject its genetic material. The *bactericidal* and *productive* ranges correspond to the phage’s ability to kill the host and to generate progeny virions, respectively. Phage-host interactions may shift along this continuum, for instance, through point mutations in critical viral proteins or changes in bacterial CRISPR spacer sequences. Recognizing this complexity of definition of host range it should be stressed that in the literature typically the term host range is understood as bactericidal range according to Hyman and in this article we are also using a term host range in such meaning.

Host range may be measured or predicted using several different approaches, which can generally be divided into three categories, each offering distinct advantages and limitations. The first category includes methods based on the observation of macroscopic features associated with ongoing bacterial lysis or growth inhibition—such as plaque formation in bacterial lawns on agar plates or changes in turbidity in liquid cultures. These assays are typically the most practical for screening studies and are the most commonly used overall [7]. The second category comprises various molecular techniques, such as fluorescent staining or hybridization of phage DNA [8, 9], flow cytometry [9], analysis of phage adsorption [10], qPCR-based approaches [11, 12], among others. These methods are usually more complex and require more advanced equipment; however, they provide detailed insights into the mechanisms of phage-host interactions. Finally, the third category includes bioinformatic approaches, such as the analysis of sequence similarities, which can be used to preliminarily select subsets of promising phage-host pairs from larger collections [13].

Despite this expanding methodological landscape, a major challenge remains: the lack of standardization across assays and experimental conditions, which hinders comparison between studies and complicates regulatory approval [14, 15, 16]. Even within culture-based methods, different authors propose distinct experimental designs and approaches to data analysis. For example, in many studies, the simplest agar-based technique is the *spot test*, in which a single, highly concentrated phage sample is applied onto soft agar containing the target bacterial host. A more precise, albeit more labor-intensive, method is the *efficiency of plating* (EOP), which involves counting plaques after the phage sample is uniformly spread on the agar surface. In the EOP approach, results are typically normalized against a reference (model) host of the tested phage [17, 18]. In assays measuring bacterial growth kinetics in liquid culture (usually spectrophotometric tests), also different endpoint metrics are used. Rajnovic et al. proposed the “percent of inhibition” [19], which is mathematically identical to the “liquid assay score” described by Xie et al. [20]; both refer to the ratio between the areas under the absorbance-vs-time curves for bacterial cultures with and without phage treatment. Konopacki et al. proposed a different calculation method, termed *PhageScore*, which also incorporates the multiplicity of infection (MOI), i.e., the ratio of phage particles to host cells, alongside a more advanced mathematical approach to curve fitting [21]. Storms et al. introduced a similar concept, the *Virulence Index*, which also accounts for MOI and additionally proposes a new standardized metric, *MV50*, defined as the MOI at which a phage achieves 50% of its maximum theoretical virulence [22]. A distinct approach is represented by the developers of *PHIDA*, an automated tool for analyzing spectrophotometric data. PHIDA can detect various types of phage-induced lysis, filter outliers and anomalies, and generate comprehensive reports. The tool is built upon a range of statistical and logical functions designed to streamline data analysis from kinetic growth inhibition experiments [23].

Agar-based and liquid culture-based methods usually show reasonably good agreement [20, 24, 25]. The main arguments against spot tests include their time-consuming nature and limited sensitivity, particularly when the lytic effect is slow or restricted to a subpopulation of host cells. Additionally, some bacterial strains are not amenable to growth on solid agar, and certain phages do not spread efficiently within the agar matrix. On the other hand, cytostatic effects observed in liquid assays do not necessarily indicate effective lysis or the production of progeny phage particles. Moreover, the rapid emergence of resistant bacterial mutants in liquid culture may lead to false-negative results [16]. Both methods, when performed at high multiplicity of infection (MOI), are susceptible to false-positive results due to a phenomenon known as *external lysis* or *lysis from without*. In such cases, large numbers of phages interacting with bacterial cell walls, or high concentrations of phage-derived lytic enzymes (e.g., endolysins, holins) present in the medium, may induce bacterial lysis that is not a consequence of the phage replication cycle [25, 26].

Therefore, a comprehensive, multi-method approach might improve accuracy of assessment of phage host range and lytic efficacy. We implemented such multi-method evaluation strategy to assess the host range and lytic activity of BAFASAL^®^, a commercial cocktail composed of four anti-*Salmonella* bacteriophages, designed to reduce *Salmonella enterica* serovar Enteritidis contamination in industrial poultry production.

*Salmonella* is a genus of Gram-negative, facultatively aerobic bacteria and is one of the leading causes of foodborne human diseases worldwide. *Salmonella enterica* serovar Enteritidis and *Salmonella enterica* serovar Typhimurium are responsible for the majority of human salmonellosis cases [27, 28]. Both serovars belong to the non-typhoidal *Salmonella* (NTS) group. It is estimated that NTS are responsible for an alarming 93 million gastrointestinal infections and approximately 155,000 diarrheal deaths annually [28, 29]. In 2022, in the European Union, *S. Enteritidis* accounted for the highest proportion (48.9%) of reported salmonellosis cases [30]. Poultry is the primary source of foodborne salmonellosis in humans and serves as the principal reservoir for *S. Enteritidis* [3, 27, 31]. In North America and Europe, *S. Enteritidis* is the dominant serovar involved in egg-borne transmission of *Salmonella* to humans [27, 30]. More than 20% of *S. Enteritidis* isolates from human cases in the EU exhibited multidrug resistance (MDR) [30]. Globally, the prevalence of MDR among NTS strains exceeds 50% in some regions of sub-Saharan Africa and Asia. Particularly alarming is the situation in Bangladesh, where 94% of *Salmonella* strains isolated from broiler chickens have been reported as multidrug-resistant [28].

Bacteriophages have been proposed as a potential solution for controlling *Salmonella* contamination, including MDR variants of *Salmonella*, in the food industry, particularly in poultry farming [3, 27, 32]. One such product is BAFASAL^®^, developed and manufactured by Proteon Pharmaceuticals S.A. (Łódź, Poland). All phage strains in BAFASAL^®^ are strictly lytic and belong to the class *Caudoviricetes* (formerly order *Caudovirales*). The preparation has demonstrated significant potential, showing favorable safety profile and high efficacy in reducing *Salmonella* levels in infected avian populations [33].

In this study, we tested BAFASAL^®^ activity against a collection of 72 unique *Salmonella* strains, including 55 *S. Enteritidis* isolates. To ensure robustness, we employed both solid media-based spot assays with serial dilutions and liquid culture-based spectrophotometric assays. Furthermore, we developed a methodology for quantitation of results of diluted spot assay and for generation of synthetic measurement of lytic activity observed in both experimental assays (Combined Lytic Score), enhancing their interpretability and predictive power. Bioinformatic analysis of bacterial genomes and antibiotic resistance profiling further contextualized the observed phage-bacteria interactions. By combining the strengths and limitations of each method, this study aims not only to investigate specificity and efficacy of BAFASAL^®^, but also to contribute to the refinement of host range determination protocols, ultimately supporting the processes of development and validation of phage-based interventions in public health, agriculture, and beyond.

## MATERIALS AND METHODS

### Collection of *Salmonella* strains

In this study, collection of tested *Salmonella enterica* strains was obtained and possessed by Proteon Pharmaceuticals. The tested collection consisted of 72 unique *S. enterica* strains, specifically: 55 specimens of serovar Enteritidis, 7 specimens of serovar Typhimurium, 3 specimens of serovar Infantis, 3 specimens of serovar Virchow and 4 of the other serovars.

All strains were cultured in LB medium (10 g/L trypton, 5 g/L yeast extract, 10 g/L NaCl, pH 7.0) for 20 hours at 37°C.

For detailed characterization the genetic material of the used strains was sequenced by NGS paired-end 2x 150 bp sequencing on Illumina MiniSeq, NovaSeq 6000 platform or ONT/Illumina sequencing. The datasets generated and analysed during the current study are available in the NCBI repository, BioProject no PRJNA1199308.

### Bioinformatic analysis

Genomic sequences of collected strains were analysed by set of bioinformatic tools. For raw sequencing data, processing and preliminary analyses were performed using the Tormes 1.3.0 [34] pipeline.

In order to confirm the classification of the analysed isolates to the particular genus, the Barrnap tool (BAsic Rapid Ribosomal RNA Predictor; T. Seemann) was used to extract 16S rRNA genes from the genomic sequences, which were then used for taxonomic classification using the RDP Classifier tool (confidence level 0.8; [35]). Further species identification was performed by the Kraken2 tool (2.1.1). *In silico Salmonella* serotype prediction was performed with SISTR tool (*Salmonella In Silico* Typing Resource; [36]) and compared with SeqSero 1.2 tool results (<u>https://</u> cge.cbs.dtu.dk/services/seqsero/; accession 08-10.2023);

Multi-Locus Sequence Typing (MLST) was determined by applying traditional PubMLST typing schemes, additionally *Salmonella* specific core genome typing was performed by means of cgMLSTFinder 1.2 tool (https://cge.food.dtu.dk/services/cgMLSTFinder/; accession 11.2023), which employs cgMLST v2 (Enterobase) scheme consisting of alleles from 3002 loci. Based on the allele profiles obtained in core genome MLST (cgMLST) analysis, the pairwise distances between genomes were established in R 4.2.1 using „gower” distance metrics (R Core Team, 2022; package stats version 4.2.1, cluster version 2.1.3). cgMLST based tree was generated from distance matrix for all 55 *Salmonella enterica* ser. Enteritidis strains employing hierarchical clustering in R (package ape version 5.6-2) and visualized in iTol v6 tool (https://itol.embl.de/; online accession 11.2023).

The presence of antibiotic resistance genes was determined by AMRFinderPlus 3.10.18 tool with the identification threshold of 60% coverage, 90% identity [37], whereas identification of virulence factors was done by VFanalyzer online platform, based on Virulence Factor DataBase (VFDB; accession: 08.-10.2023) [38, 39]. *Salmonella* Pathogenicity Islands were identified by means of SPIFinder-2.0 tool (https://cge.cbs.dtu.dk/services/SPIFinder/; accession: 09.-10.2023), provided by the Center for Genomic Epidemiology.

Sequences were also analysed for the presence of prophage insertions using online PHASTEST tool (https://phastest.ca; [40, 41]).

For the purpose of whole-genome diversification the strains were assessed for average nucleotide identity (ANI) (pyani v. 0.2.10; [42]).

### Antibiotic susceptibility assay

Susceptibility of all *Salmonella* strains to Sulphamethoxazole/Trimethoprim (Oxoid), Enrofloxacin (Oxoid), Gentamycin (Oxoid) and Ampicillin (Oxoid) was determined using disk diffusion method. To this end, 90 mm Mueller Hinton agar plates were inoculated by spreading evenly bacterial suspensions of OD_600_=0.5 with cotton swab. Subsequently, plates were divided into 6 zones, and within each zone, disk impregnated with a known concentration of antibiotic from disk dispenser was attached to the agar plate with the use of sterile forceps. Subsequently, test plates were incubated at 37°C for 24 hours and results were interpreted based on growth inhibition zone diameter, classified in accordance with Eucast Clinical Breakpoints table version 13.1 valid from 29.08.2023 except for the results for Enrofloxacin, classified in accordance to CLSI VET 01S-ED6-2023 (update 20.02.2023) as being susceptible, intermediate or resistant to an antimicrobial agent.

### Bacteriophage preparation

BAFASAL^®^ is a cocktail of four anti-*Salmonella* bacteriophages, deposited in the Polish Collection of Microorganisms under accession numbers: PCM F/00069 (strain 8 sent 1748), PCM F/00070 (strain 8 sent 65), PCM F/00071 (strain 3 sent 1), and PCM F/0097 (strain 5 sent 1). For the purpose of this study, the preparation batch no. PP22B003 was manufactured with a titer 1.30x10^8^ PFU/mL as described in Wojcik et al. [33]. Briefly, each bacteriophage included in BAFASAL^®^ preparation was amplified in a separate culture. In this procedure, the culture medium was inoculated with the host production strain and incubated at 37°C for approximately 2 h. Next, the selected phage was added to the bacterial culture, and incubation proceeded for a further 3 h at 37°C leading to cell lysis. Upon amplification, the biomass was separated from the phage-containing culture fluid by filtration. Once the amplification procedure was completed, the titer of the bacteriophage was assessed by phage enumeration using a double agar overlay plaque assay according to Kropinski et al. [43] and calculated for about 1x10^8^ PFU/mL in a final mixture completed with sterile water. BAFASAL^®^ was finally tested by phage enumeration using the double agar overlay plaque assay, analyzing the microbiological sterility of the product and the presence of bacterial genomic residues according to EFSA’s “Guidance on the characterization of microorganisms used as feed additives or as production organisms.” [44]

### Serial dilutions spot test

Phage susceptibility tested by serial dilutions spot test was performed according to Kutter [17]. Chosen *Salmonella* strains were cultured in liquid LB medium until optical density of OD_600_=0.5 was reached. Next, LB agar plates were covered with top layer of the 6 ml of molten, chilled to 50°C LB agar (0.7%), with addition of 100 µl of *Salmonella* culture and left for solidification. Phage preparation was serially diluted in bacteria growth medium. Each step was a 10-fold dilution from the original concentration 10^0^ (E0) to the most diluted sample of 10^-5^ (E-5). On the surface of each plate, 10 µl of each phage dilution along with empty medium as negative control was added as separate spots. Subsequently, plates were incubated for 24 h at 37°C. After the incubation, each plate was assessed visually and results were noted, using following decreasing scale, describing intensity of the observed lysis: CL – “clear spot” (no signs of bacterial growth), T – “turbid spot” (turbidity observed on the clearly brighter spot), P – “plaques” (plaques observed in the bacterial lawn), NL – “no clearance in the spot” (no signs of the bacterial lysis observed). Each *Salmonella* strain was tested in 3 replicates started from 3 independent bacterial cultures each. Final result of the test was interpreted as follows: any lytic event (CL, T, P) was considered positive. If in 3 tested replicates, at least 2 positive events were observed in the more diluted half of the samples (from E-3 to E-5), tested strain was considered sensitive. Else, if at least 2 positive events were observed in less diluted half of the samples (from E0 to E-2), the strain was considered medium sensitive. Else, the strain was considered insensitive.

### Transformation of the serial dilutions spot test results to a continuous scale

Obtained serial dilutions spot test patterns were transformed to numerical values using MPN method to calculate the most probable number of phages from BAFASAL^®^ required to observe visible lysis on specific bacterial strain. For this purpose, the results of susceptibility testing by serial dilutions spot test from 3 biological replicates were presented in the form of the compact configuration of the number of positive events (of 3 in total) in a series of subsequent dilutions, i.e. (E0, E-1, E-2, E-3, E-4, E-5). MPN values were calculated with the use of the online calculator delivered by U.S. Environmental Protection Agency EPA (https://mostprobablenumbercalculator.epa.gov/mpnForm), with bias (Salama/Thomas methods) corrected central values and 95% confidence intervals calculated with use of the Cornish and Fisher method (max of 30 iterations). For samples, in which no lytic activity was observed in all concentrations, a censored value equal to the half of the minimum concentration possible to be determined was chosen, i.e. for configuration (1,0,0,0,0,0): MPN = 30.4 (5.0÷122.8) [central value and 95% CI] and for configuration (0,0,0,0,0,0) censored value is chosen to be 15.2 (2.5÷61.4). For samples, in which lytic activity was observed in all concentrations, a censored value equal to the twice of the maximum concentration possible to be determined was chosen, i.e. for configuration (3,3,3,3,3,2) MPN = 1.10E7 (2.25E6÷3.08E7) and for configuration (3,3,3,3,3,3) censored value is chosen to be 2.20E7 (4.50E6÷6.15E7). For each sample two MPN values were calculated. First one, (MPN_1_) describing total lytic effect, in which case all positive observations (CL, T and P) were treated as 1 and negative observations (NL) as 0 and the second one (MPN_2_) describing only strong lytic effect in which case only CL events were treated as 1 and all the rest (T, P and NL) as 0. Finally, Effective Concentration Threshold (ECT) and serial dilutions spot test score were evaluated. As absolute amounts of bacteriophages used in each experiment were known and equal across all replications, thus the differences in calculated MPN values were assumed to depend on individual susceptibility or resistance shown by different bacterial strains. To describe that value quantitatively and take into account concentration of phages in tested preparation (to standardize future results from different phage preparations) a new metric of Effective Concentration Threshold has been proposed. ECT refers to the phages concentration in tested preparation divided by MPN value (first or second) for given bacterial strain and is read as “minimal concentration of phages required to obtain a noticeable lytic effect”. Score for the serial dilutions spot test was calculated as a negative sum of the logarithms of the ECT1 and ECT2, according to the formula below in which: C_F_ – concentration of bacteriophages in undiluted preparation [PFU/mL]; MPN_1_, MPN_2_ – calculated concentrations according to total lytic effect and strong lytic effect respectively [MPN/mL]. Minus sign was added to keep directly proportional relationship between test score value and susceptibility of the tested strain.

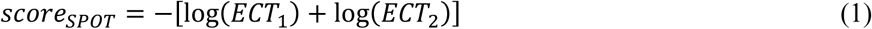

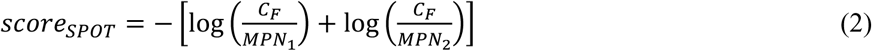

### Growth inhibition test

Spectrophotometric, kinetic growth inhibition test was performed according to Xie et al. [20]. *Salmonella* strains were cultured in liquid LB medium until optical density of OD_600_=0.5 was reached. Test was performed on a standard, sterile, clear, flat-bottom 96-well plates. Each tested bacterial culture was dispensed into 6 wells, 100 µl per well. Subsequently, to 3 of the wells 100 µl of phage preparation was added and to the other 3 wells: 100 µl of empty phage buffer. Additionally, following control conditions were tested on each plate, each in 3 replicates: medium control of 100 µl of empty medium and 100 µl of empty buffer; phage control of 100 µl of empty medium and 100 µl of phage preparation. Assembled plates were subsequently placed in a plate reader and incubated for 8 hours with gentle shaking and chamber temperature set at 37°C. Every 20 minutes absorbance at 620 nm was measured.

### Data analysis from growth inhibition test

All results from reader were first carefully assessed. 9 of 243 technical replicates of control wells were excluded from analysis due to observed bacterial growth (in no bacteria controls). Furthermore, any abnormal, strong outlying results (e.g. caused by air bubble in a well) was removed and replaced with a mean value from residual replicates (less than 0.06% of the total collected data points from control and experimental wells was treated this way).

Next, for each plate, for each time point blank value was calculated by averaging reads from 3 control wells. Subsequently, proper blank values were subtracted from the reads from experimental samples. To compare bacterial growth, area under optical density curve (AUC) over time was calculated independently for each well. AUC was calculated according to the formula below (for n time points), where OD refers to the optical density value at the given time point.

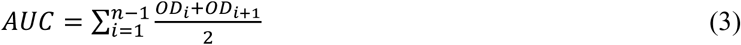

Subsequently, calculated AUC values from 3 technical replicates were tested for possibility of outliers with Grubbs’ test (α=0.05) and if outlier was detected, such replication was rejected. Remaining values were averaged, and AUC normalization score (ANS) was calculated according to the formula below, where AUC_control_ is mean AUC without phages and AUC_phages_ is mean AUC in presence of bacteriophages, for given bacterial strain.

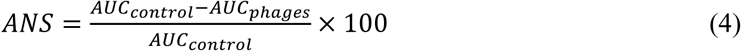

Formulas for AUC and ANS were taken directly from the work of Xie et al. [20]. Measurements for each *Salmonella* strain were replicated 3 times in 3 independent experiments. Subsequently, obtained ANS values were also tested with Grubbs’ test, and if it was justified, outlying value was rejected. Final result is presented as mean±SD from 2 or 3 biological replicates. At the end, the tested strain is interpreted as sensitive if mean ANS is greater than 15% and insensitive in the opposing case.

### Normalization and calculation of the Combined Lytic Score

Results from the serial dilutions spot test and spectrophotometric analysis of kinetics of bacterial growth test were normalized using top-bottom method to the range of 0-100%. Normalized values were calculated with use of the equation 5 below, where value_min_ and value_max_ are presented in **Table 1**.

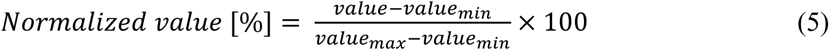

**Table 1.**
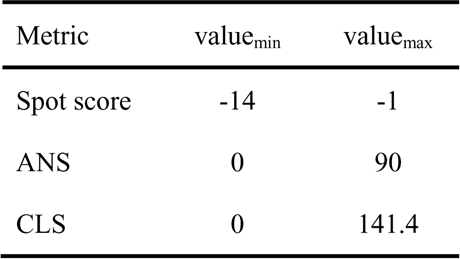
Coefficients used in data normalization

Combined Lytic Score (CLS) values were calculated according to equation 6 below, where “norm. spot score” and “norm. ANS” are normalized results of the partial tests (serial dilutions spot test and spectrophotometric analysis of kinetics of bacterial growth test respectively) and weights (0.725 and 0.689) are coordinates of the unit vector parallel to the regression line fitted to the calibration dataset:

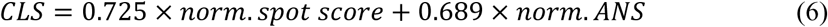

Next, CLS values were normalized to the percentage scale. Additionally, all preceding equations were algebraically combined to obtain a single simplified formula (Eq. 7) for convenient calculation of normalized CLS values.

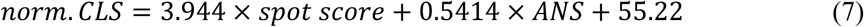

### Statistical Analysis

Data are presented as mean ± SD. Outlying points in screening were detected with Grubbs’ test with (α=0.05). Screening data pipelining was conducted in Python. Significance of differences in CLS scores between compared groups was assessed with Mann-Whitney test, with use of StatsDirect 4.0.4 software.

## RESULTS

### *In vitro* phage susceptibility assays

72 strains of *Salmonella enterica* were tested in 3 biological replicates for susceptibility towards phage lysis, using the same BAFASAL^®^ preparation as phage source in each experiment. Results are presented in **Table 2**. Partial results of serial dilutions spot test are presented as number of positive lysis observations in the series of subsequent phage dilutions and results of kinetic growth inhibition (ANS) describes how much area under growth curve was inhibited in presence of phages.

**Table 2.** Results of the *in vitro* susceptibility tests: S – susceptible, medS – medium susceptible, INS – insusceptible.

| Strain | Spot test |  |  | Growth inhibition test |  |  |
| --- | --- | --- | --- | --- | --- | --- |
|  | Any lytic | CL only | Result | mean ANS | SD | Result |
| Brandenburg 584 | 3,3,3,3,0,0 | 0,0,0,0,0,0 | S | 13,71 | 0,09 | INS |
|  | Any lytic | CL only | Result | mean ANS | SD | Result |
| Enteritidis 012PP2014 | 3,3,3,3,3,3 | 3,3,3,3,0,0 | S | 70,67 | 3,77 | S |
| Enteritidis 070PP2018 | 3,3,3,3,3,3 | 3,3,3,1,1,0 | S | 69,97 | 2,13 | S |
| Enteritidis 071PP2018 | 3,3,3,3,3,3 | 3,3,3,2,0,0 | S | 67,17 | 3,45 | S |
| Enteritidis 093PP2018 | 3,3,3,3,3,3 | 3,3,3,3,0,0 | S | 77,06 | 2,70 | S |
| Enteritidis 1013/S/09 K4 | 3,3,3,3,3,3 | 3,3,3,3,3,2 | S | 79,71 | 3,12 | S |
| Enteritidis 1014/S/09 K1 | 3,3,3,3,3,3 | 3,3,3,3,3,2 | S | 66,12 | 15,28 | S |
| Enteritidis 1044/S/09 K2 | 3,3,3,3,3,3 | 3,3,3,3,3,2 | S | 84,65 | 4,55 | S |
| Enteritidis 1047/S/09 K1 | 3,3,3,3,3,3 | 3,3,3,3,3,2 | S | 76,81 | 6,41 | S |
| Enteritidis 1061/S/09 K5 | 3,3,3,3,3,3 | 3,3,3,3,3,3 | S | 70,21 | 3,29 | S |
| Enteritidis 1067/S/09 K2 | 3,3,3,3,3,3 | 3,3,3,3,3,3 | S | 79,91 | 4,62 | S |
| Enteritidis 1085/S/09 MEK | 3,3,3,3,3,3 | 3,3,3,3,3,3 | S | 80,67 | 0,40 | S |
| Enteritidis 114PP2019 | 3,3,3,3,3,3 | 3,3,3,3,0,0 | S | 75,44 | 6,38 | S |
| Enteritidis 1171/S/09 | 3,3,3,3,3,3 | 0,0,0,0,0,0 | S | 58,21 | 9,97 | S |
| Enteritidis 1250/S/09 | 3,3,3,3,3,3 | 0,0,0,0,0,0 | S | 68,66 | 9,91 | S |
| Enteritidis 1445/S/09 K46 | 3,3,3,3,3,2 | 0,0,0,0,0,0 | S | 35,29 | 23,04 | S |
| Enteritidis 1446/S/09 K31 | 3,3,3,3,3,3 | 2,0,0,0,0,0 | S | 38,88 | 3,06 | S |
| Enteritidis 1535/S/09 | 3,3,3,3,3,3 | 0,0,0,0,0,0 | S | 53,28 | 0,38 | S |
| Enteritidis 1545/S/09 MEK | 3,3,3,3,3,3 | 3,3,3,3,3,3 | S | 79,29 | 1,04 | S |
| Enteritidis 1573/S/09 NWJ | 3,3,3,3,3,3 | 3,3,3,3,3,0 | S | 80,21 | 2,84 | S |
| Enteritidis 1714/09 | 3,3,3,3,3,3 | 3,0,0,0,0,0 | S | 79,20 | 0,94 | S |
| Enteritidis 1748 | 3,3,3,3,3,3 | 3,3,3,3,2,2 | S | 75,40 | 4,57 | S |
| Enteritidis 2050/S/09 K4 nr2 | 3,3,3,3,3,3 | 0,0,0,0,0,0 | S | 38,59 | 0,59 | S |
| Enteritidis 212PP2021 | 3,3,3,2,2,2 | 1,1,1,1,0,0 | S | 50,65 | 12,86 | S |
| Enteritidis 249 | 3,3,3,3,3,3 | 3,3,3,3,3,3 | S | 81,40 | 3,30 | S |
| Enteritidis 423PP2021 | 3,3,3,3,3,3 | 3,3,3,2,0,0 | S | 50,31 | 1,03 | S |
| Enteritidis 455PP2021 | 3,3,3,3,3,3 | 3,3,3,1,0,0 | S | 47,79 | 16,49 | S |
| Enteritidis 462PP2021 | 3,3,3,3,3,3 | 3,3,3,3,0,0 | S | 67,31 | 11,73 | S |
| Enteritidis 465PP2022 | 3,3,3,3,3,3 | 3,3,3,3,0,0 | S | 80,04 | 0,64 | S |
| Enteritidis 466PP2022 | 3,3,3,3,3,3 | 3,3,3,3,0,0 | S | 71,85 | 7,77 | S |
| Enteritidis 470PP2022 | 3,3,3,3,3,3 | 3,3,3,3,0,0 | S | 63,25 | 0,45 | S |
| Enteritidis 472PP2022 | 3,3,3,3,3,3 | 3,3,3,1,0,0 | S | 77,96 | 4,17 | S |
| Enteritidis 476PP2022 | 3,3,3,3,3,3 | 3,3,3,0,0,0 | S | 29,30 | 3,74 | S |
| Enteritidis 477PP2023 | 3,3,3,3,3,3 | 3,3,3,3,0,0 | S | 64,69 | 1,73 | S |
| Enteritidis 488PP2023 | 3,3,3,3,3,3 | 1,1,0,0,0,0 | S | 11,84 | 0,54 | INS |
| Enteritidis 489PP2023 | 3,3,3,3,3,3 | 3,3,3,3,0,0 | S | 80,23 | 1,52 | S |
| Enteritidis 490PP2023 | 3,3,3,3,3,3 | 3,3,3,3,0,0 | S | 79,16 | 7,54 | S |
| Enteritidis 491PP2023 | 3,3,3,3,3,3 | 3,3,3,3,0,0 | S | 77,00 | 5,19 | S |
| Enteritidis 492PP2023 | 3,3,3,3,3,3 | 3,3,3,3,0,0 | S | 75,61 | 3,96 | S |
| Enteritidis 493PP2023 | 3,3,3,3,3,3 | 3,3,3,3,0,0 | S | 63,85 | 14,91 | S |
| Enteritidis 494PP2023 | 3,3,3,3,3,3 | 3,3,3,3,0,0 | S | 51,82 | 0,25 | S |
| Enteritidis 517/S/09 | 3,3,3,3,3,3 | 3,3,3,3,3,2 | S | 86,64 | 0,73 | S |
| Enteritidis 571/S/09 | 3,3,3,3,3,3 | 3,3,3,3,3,1 | S | 68,42 | 9,91 | S |
| Enteritidis 64/S/10 | 3,3,3,3,3,3 | 3,3,3,3,3,2 | S | 79,29 | 4,93 | S |
| Enteritidis 65/S/10 | 3,3,3,3,3,3 | 3,3,3,3,1,0 | S | 65,56 | 5,85 | S |
| Enteritidis 833/S/09 | 3,3,3,3,3,3 | 3,3,3,3,3,2 | S | 78,35 | 4,27 | S |
|  | Any lytic | CL only | Result | mean ANS | SD | Result |
| Enteritidis 838/S/09 B | 3,3,3,3,3,3 | 0,0,0,0,0,0 | S | 38,28 | 0,97 | S |
| Enteritidis 866/S/09 | 3,3,3,3,3,3 | 3,3,3,3,3,2 | S | 75,98 | 4,21 | S |
| Enteritidis 945/S/09 | 3,3,3,3,3,3 | 2,0,0,0,0,0 | S | 68,96 | 1,90 | S |
| Enteritidis D ATCC 13076 | 3,3,3,3,3,3 | 3,3,3,3,3,1 | S | 68,44 | 6,51 | S |
| Enteritidis KK10 | 3,0,0,0,0,0 | 0,0,0,0,0,0 | med.S | 22,36 | 1,08 | S |
| Enteritidis KK11 | 1,0,0,0,0,0 | 0,0,0,0,0,0 | INS | 14,07 | 14,19 | INS |
| Enteritidis KK3 | 3,3,3,3,3,3 | 3,3,3,3,3,3 | S | 83,08 | 5,56 | S |
| Enteritidis KK6 | 3,3,3,3,3,3 | 3,3,3,3,3,0 | S | 75,26 | 3,89 | S |
| Enteritidis KK9 | 3,3,0,0,0,0 | 0,0,0,0,0,0 | med.S | 17,80 | 0,11 | S |
| Enteritidis PK1 | 3,3,3,3,3,3 | 3,3,3,3,3,1 | S | 78,70 | 8,01 | S |
| Gallinarum 001PP2015 | 3,3,3,3,3,3 | 3,3,3,3,0,0 | S | 68,82 | 26,24 | S |
| Hadar 817 | 3,3,3,3,3,2 | 0,0,0,0,0,0 | S | 14,09 | 11,72 | INS |
| Infantis 1021/S/09 | 0,0,0,0,0,0 | 0,0,0,0,0,0 | INS | -2,00 | 0,18 | INS |
| Infantis 144 | 0,0,0,0,0,0 | 0,0,0,0,0,0 | INS | -0,38 | 0,60 | INS |
| Infantis ZK5 | 3,3,3,1,0,0 | 0,0,0,0,0,0 | med.S | -0,51 | 0,10 | INS |
| Paratyphi A ATCC 19150 | 3,3,3,2,0,0 | 0,0,0,0,0,0 | S | 18,29 | 3,87 | S |
| Typhimurium 1081/S/09 K1 | 3,3,0,0,0,0 | 0,0,0,0,0,0 | med.S | 6,51 | 0,04 | INS |
| Typhimurium 1567/S/09 | 3,3,3,3,3,0 | 0,0,0,0,0,0 | S | 16,09 | 3,02 | S |
| Typhimurium 1578/S/09 | 3,0,0,0,0,0 | 0,0,0,0,0,0 | med.S | 10,84 | 0,13 | INS |
| Typhimurium 1785/09 | 0,0,0,0,0,0 | 0,0,0,0,0,0 | INS | 7,52 | 1,65 | INS |
| Typhimurium 531 | 2,0,0,0,0,0 | 0,0,0,0,0,0 | med.S | 3,23 | 0,22 | INS |
| Typhimurium 960/S/09 | 3,3,3,3,3,2 | 0,0,0,0,0,0 | S | 4,13 | 9,20 | INS |
| Typhimurium ATCC 13311 | 3,3,3,3,3,3 | 0,0,0,0,0,0 | S | 35,71 | 3,84 | S |
| Virchow 1175/S/09 | 0,0,0,0,0,0 | 0,0,0,0,0,0 | INS | -2,00 | 0,75 | INS |
| Virchow 1188/S/09 K10 | 0,0,0,0,0,0 | 0,0,0,0,0,0 | INS | 2,48 | 2,91 | INS |
| Virchow GAP K2 | 0,0,0,0,0,0 | 0,0,0,0,0,0 | INS | -3,74 | 0,56 | INS |

### Analysis of the correlation between two lytic activity assays

Summary of results from individual *in vitro* assays are presented in **Table 3** and **Table 4**. From this simple binning we can observe 86.1% agreement in results between both methods. However, it is worth mentioning that serial dilutions spot assay has 3 possible outcomes, while growth inhibition assay only 2, which induces high bias in this value.

**Table 3.** Distribution of outcomes of the *in vitro* susceptibility assays

| Result | Serial dilutions spot test | Growth inh. test |
| --- | --- | --- |
| sensitive | 59 | 57 |
| medium sensitive | 6 | n/a |
| insensitive | 7 | 15 |

**Table 4.** Agreement of results from both assays in the corresponding samples

| Outcome | n | % |
| --- | --- | --- |
| same | 62 | 86.1 |
| different | 10 | 13.9 |
| total | 72 | 100 |

To better understand correlation between the presented results, serial dilutions spot test results were transformed into numerical values on continuous scale. Data regarding positive observations (already shown in **Table 2**) from serial dilutions spot test were extracted and MPN method was utilized to calculate most probable number of (apparent) phages in each strain, subsequently pot test score was calculated, according to equation 1 and 2 presented in Materials and Methods.

At first the correlation using only the total lytic effect from the serial dilutions spot test (where all visible signs of lysis were treated as positive outcome or in other words, the ECT2 component in serial dilutions spot test score formula had zero weight) was analyzed. Scatter plot with linear regression line is presented in **Figure 1A**. Despite relatively high R2 coefficient value suggesting strong correlation, a large group of aggregated points are placed at the right end of the chart (points with maximum serial dilutions spot test score, which suggests that this metric is not optimal for this experimental model).

**Figure 1.**
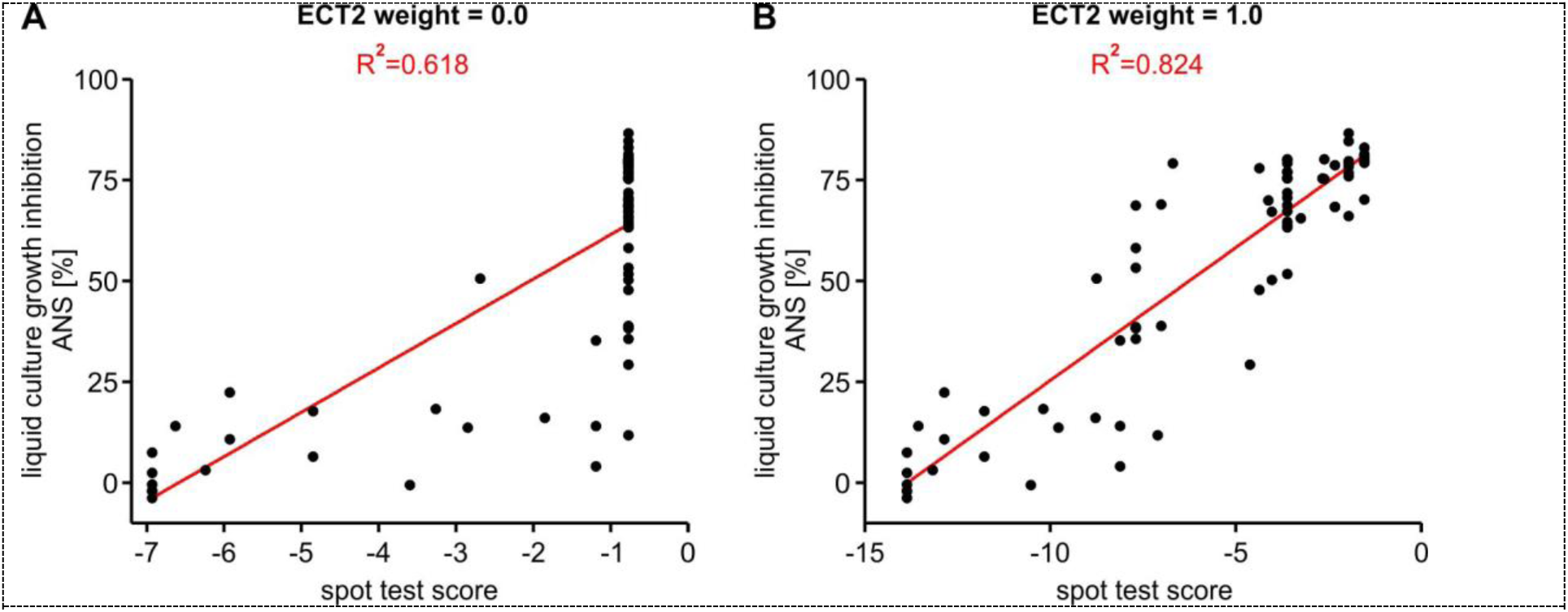
Correlation of the results from two phage susceptibility *in vitro* assays. Red: linear regression line fitted to data. A: using simplified formula for the serial dilutions spot test, considering only total lytic component; B: using optimized formula for serial dilutions spot test, total lytic component and strong lytic component of equal weights.

To solve this problem, we tried to extract more information from serial dilutions spot test raw data adding a strong lytic effect as additional category of visual observation. It was represented by additional component in equation, where only the clear lysis observations were treated as positive outcome. However, the proper weight of the second component in the equation reflecting real dependencies was unknown. For this purpose, computer optimization (Solver Add-in of the MS Excel) was used to maximize R2 value of the correlation between both tests (target function), with change of the ECT2 weight in serial dilutions spot test score equation (the changed variable). Best R2 value (0.8243) was obtained at ECT2 weight of 1.127. However, a close value of 1.0 was chosen to simplify resulting formula. With such simplification, obtained R2 (0.8236) is still very close to the Best R2. Scatter plot created with the use of the optimized formula is presented in **Figure 1B**.

### Combined Lytic Score calculation

Using the data points from correlation analysis (with optimal serial dilutions spot assay score equation) the outcome of both assays was normalized to the 0-100% range. In the first analysis we used the lowest and highest values from a given set as the normalization limit between 0% and 100%. However, although this approach may be considered appropriate for a single, large data set, as presented in this study, it would not work if one wanted to apply this method to a small number of strains. In order to obtain more objective normalization limits, we carefully analyzed the results of both *in vitro* tests. We chose limits from 0 to 90% growth inhibition for the kinetic test (since no sample, even the most susceptible to lysis, exceeds 90% inhibition). For the serial dilutions spot test, the obtained results range from -13.86 to -1.54. Since the concentration of the bacteriophage used is a component of the serial dilutions spot test result, we expect that the results obtained in other studies may exceed these limits using more or less diluted phage samples, therefore we arbitrarily chose values from -14 to -1. Then, the obtained results were normalized based on these values. It is worth noting that in the case of the spectrophotometric test, some of the normalized values go below 0, these are samples in which, on average, the strain after adding the bacteriophage grew slightly better than the control (within the margin of statistical error).

Subsequently, normalized data points were plotted on a X-Y plot. By looking on the scatter plot drawn from these points (**Figure 2A**) the region of high susceptibility (top, right corner) where both assays are close to the maximal values and region of high resistance (bottom, left corner) was easily identified. To quantify unified result from both assays with a single value, a new metric of Combined Lytic Score (CLS) which is defined as the scalar product of the point coordinates on the mentioned scatter plot with unit vector parallel to the regression line fitted to data was proposed. In other words, this metric describes how close the point is to the top-right (susceptible) corner of the scatter plot, measuring distance only in component parallel to the regression line. For convenience, CLS value is also presented after normalization (0-100% range). Normalized CLS values were calculated for all the strains tested in this study, results are presented in **Figure 2B**. 52% of the tested strains have reached normalized CLS of over 75%, all but one of them are of serovar Enteritidis. The only non-Enteritidis strain from the high susceptible region (marked with # on a figure) is Gallinarum 001PP2015, which is control strain for viability and lytic activity of BAFASAL^®^. Individual *in vitro* test scores and CLS, all raw and normalized are presented also in **Table 5**.

**Figure 2.**
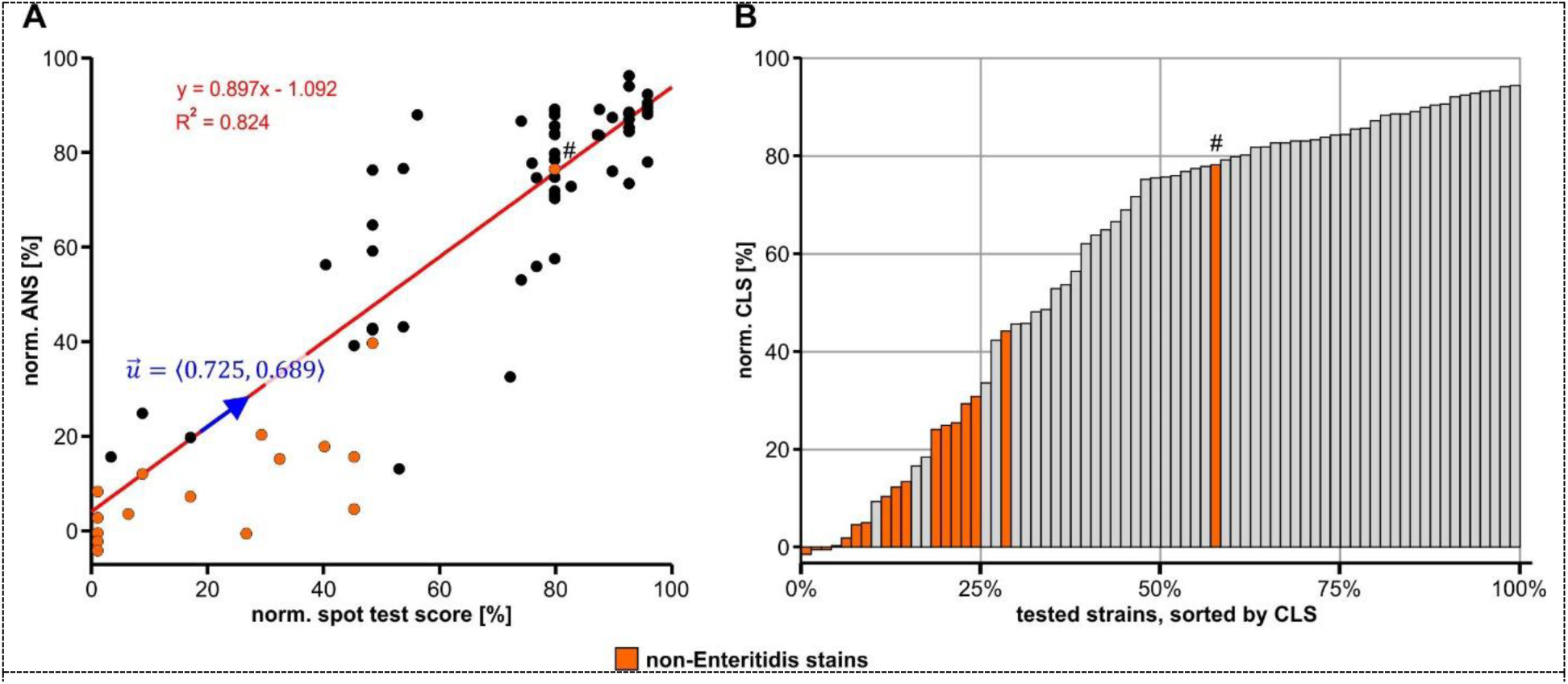
Combined Lytic Score. **A**: scatter plot of the normalized results from both assays, regression line (red) and symbolic unit vector (blue) parallel to the regression line; **B**: histogram of the CLS values, sorted in ascending order, reached by individual strains tested in this study; **#** - Gallinarum 001PP2015, control strain for BAFASAL^®^.

**Table 5.**
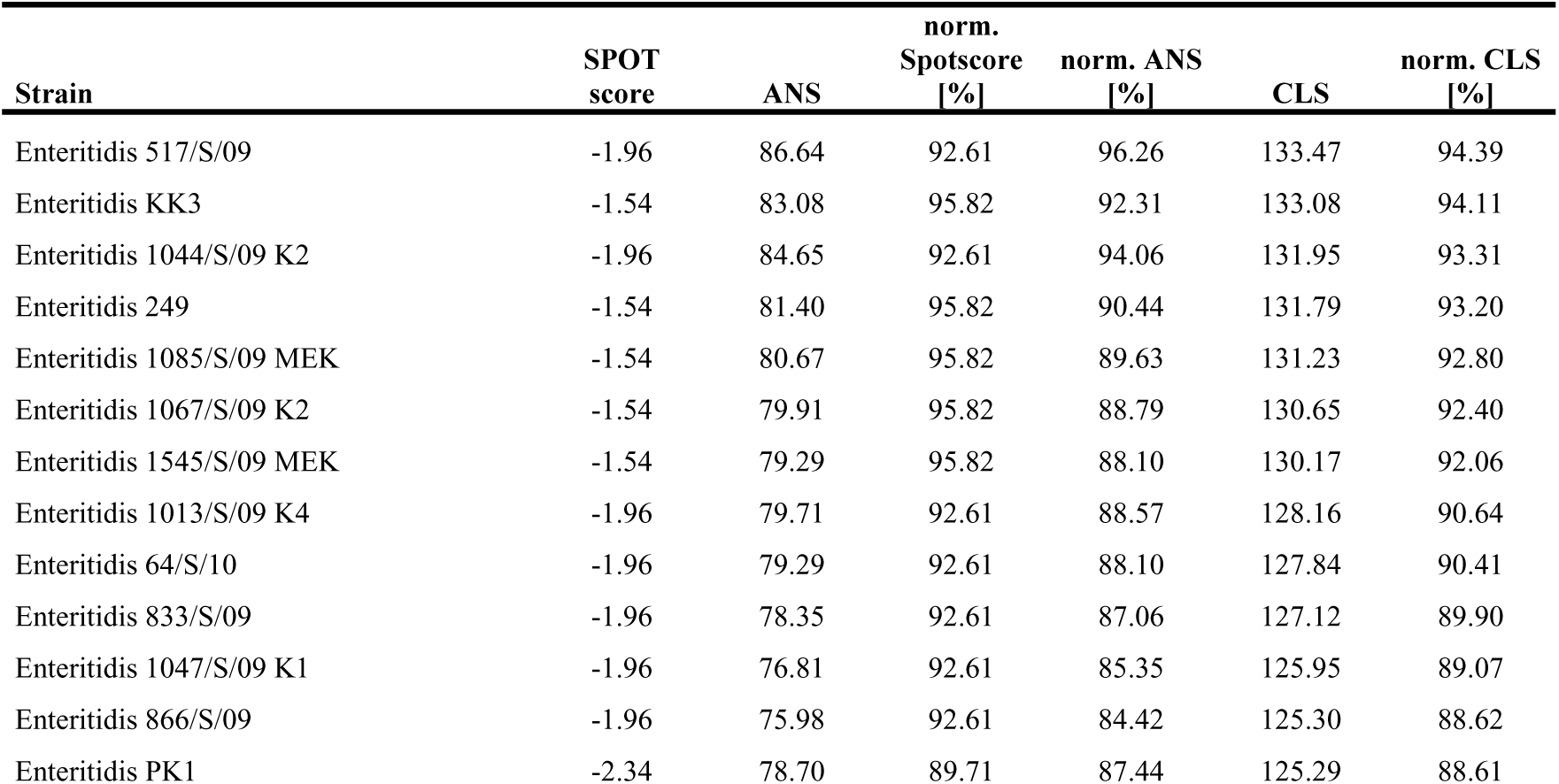
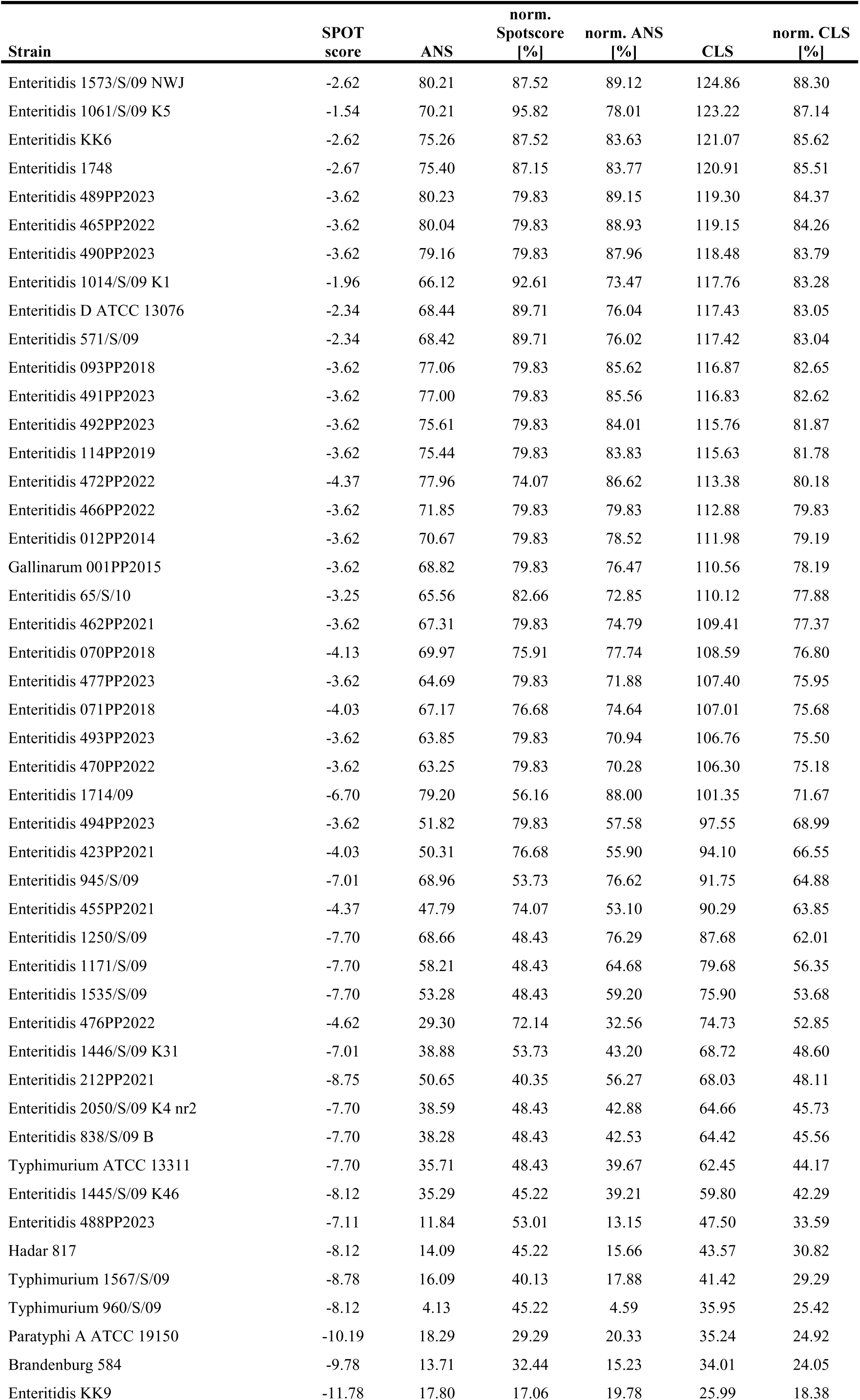
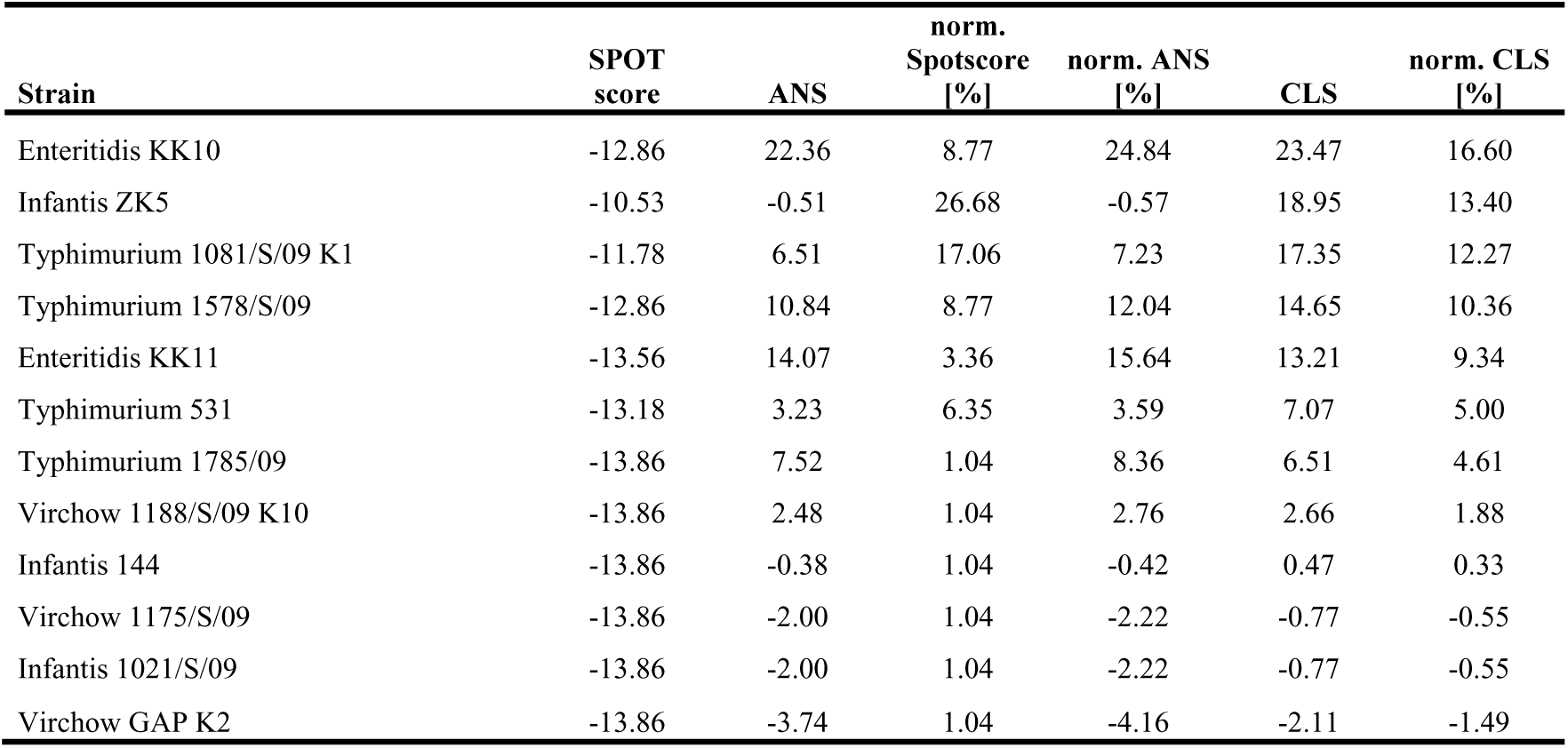
Raw and normalized results of *in vitro* susceptibility assays and Combined Lytic Score. Strains sorted according to calculated CLS value, from most susceptible to most resistant.

| Strain | SPOT score | ANS | norm. Spotscore [%] | norm. ANS [%] | CLS | norm. CLS [%] |
| --- | --- | --- | --- | --- | --- | --- |
| Enteritidis 517/S/09 | -1.96 | 86.64 | 92.61 | 96.26 | 133.47 | 94.39 |
| Enteritidis KK3 | -1.54 | 83.08 | 95.82 | 92.31 | 133.08 | 94.11 |
| Enteritidis 1044/S/09 K2 | -1.96 | 84.65 | 92.61 | 94.06 | 131.95 | 93.31 |
| Enteritidis 249 | -1.54 | 81.40 | 95.82 | 90.44 | 131.79 | 93.20 |
| Enteritidis 1085/S/09 MEK | -1.54 | 80.67 | 95.82 | 89.63 | 131.23 | 92.80 |
| Enteritidis 1067/S/09 K2 | -1.54 | 79.91 | 95.82 | 88.79 | 130.65 | 92.40 |
| Enteritidis 1545/S/09 MEK | -1.54 | 79.29 | 95.82 | 88.10 | 130.17 | 92.06 |
| Enteritidis 1013/S/09 K4 | -1.96 | 79.71 | 92.61 | 88.57 | 128.16 | 90.64 |
| Enteritidis 64/S/10 | -1.96 | 79.29 | 92.61 | 88.10 | 127.84 | 90.41 |
| Enteritidis 833/S/09 | -1.96 | 78.35 | 92.61 | 87.06 | 127.12 | 89.90 |
| Enteritidis 1047/S/09 K1 | -1.96 | 76.81 | 92.61 | 85.35 | 125.95 | 89.07 |
| Enteritidis 866/S/09 | -1.96 | 75.98 | 92.61 | 84.42 | 125.30 | 88.62 |
| Enteritidis PK1 | -2.34 | 78.70 | 89.71 | 87.44 | 125.29 | 88.61 |

| Strain | SPOT<br>score | ANS | norm.<br>Spotscore<br>[%] | norm. ANS<br>[%] | CLS | norm. CLS<br>[%] |
| --- | --- | --- | --- | --- | --- | --- |
| Enteritidis 1573/S/09 NWJ | -2.62 | 80.21 | 87.52 | 89.12 | 124.86 | 88.30 |
| Enteritidis 1061/S/09 K5 | -1.54 | 70.21 | 95.82 | 78.01 | 123.22 | 87.14 |
| Enteritidis KK6 | -2.62 | 75.26 | 87.52 | 83.63 | 121.07 | 85.62 |
| Enteritidis 1748 | -2.67 | 75.40 | 87.15 | 83.77 | 120.91 | 85.51 |
| Enteritidis 489PP2023 | -3.62 | 80.23 | 79.83 | 89.15 | 119.30 | 84.37 |
| Enteritidis 465PP2022 | -3.62 | 80.04 | 79.83 | 88.93 | 119.15 | 84.26 |
| Enteritidis 490PP2023 | -3.62 | 79.16 | 79.83 | 87.96 | 118.48 | 83.79 |
| Enteritidis 1014/S/09 K1 | -1.96 | 66.12 | 92.61 | 73.47 | 117.76 | 83.28 |
| Enteritidis D ATCC 13076 | -2.34 | 68.44 | 89.71 | 76.04 | 117.43 | 83.05 |
| Enteritidis 571/S/09 | -2.34 | 68.42 | 89.71 | 76.02 | 117.42 | 83.04 |
| Enteritidis 093PP2018 | -3.62 | 77.06 | 79.83 | 85.62 | 116.87 | 82.65 |
| Enteritidis 491PP2023 | -3.62 | 77.00 | 79.83 | 85.56 | 116.83 | 82.62 |
| Enteritidis 492PP2023 | -3.62 | 75.61 | 79.83 | 84.01 | 115.76 | 81.87 |
| Enteritidis 114PP2019 | -3.62 | 75.44 | 79.83 | 83.83 | 115.63 | 81.78 |
| Enteritidis 472PP2022 | -4.37 | 77.96 | 74.07 | 86.62 | 113.38 | 80.18 |
| Enteritidis 466PP2022 | -3.62 | 71.85 | 79.83 | 79.83 | 112.88 | 79.83 |
| Enteritidis 012PP2014 | -3.62 | 70.67 | 79.83 | 78.52 | 111.98 | 79.19 |
| Gallinarum 001PP2015 | -3.62 | 68.82 | 79.83 | 76.47 | 110.56 | 78.19 |
| Enteritidis 65/S/10 | -3.25 | 65.56 | 82.66 | 72.85 | 110.12 | 77.88 |
| Enteritidis 462PP2021 | -3.62 | 67.31 | 79.83 | 74.79 | 109.41 | 77.37 |
| Enteritidis 070PP2018 | -4.13 | 69.97 | 75.91 | 77.74 | 108.59 | 76.80 |
| Enteritidis 477PP2023 | -3.62 | 64.69 | 79.83 | 71.88 | 107.40 | 75.95 |
| Enteritidis 071PP2018 | -4.03 | 67.17 | 76.68 | 74.64 | 107.01 | 75.68 |
| Enteritidis 493PP2023 | -3.62 | 63.85 | 79.83 | 70.94 | 106.76 | 75.50 |
| Enteritidis 470PP2022 | -3.62 | 63.25 | 79.83 | 70.28 | 106.30 | 75.18 |
| Enteritidis 1714/09 | -6.70 | 79.20 | 56.16 | 88.00 | 101.35 | 71.67 |
| Enteritidis 494PP2023 | -3.62 | 51.82 | 79.83 | 57.58 | 97.55 | 68.99 |
| Enteritidis 423PP2021 | -4.03 | 50.31 | 76.68 | 55.90 | 94.10 | 66.55 |
| Enteritidis 945/S/09 | -7.01 | 68.96 | 53.73 | 76.62 | 91.75 | 64.88 |
| Enteritidis 455PP2021 | -4.37 | 47.79 | 74.07 | 53.10 | 90.29 | 63.85 |
| Enteritidis 1250/S/09 | -7.70 | 68.66 | 48.43 | 76.29 | 87.68 | 62.01 |
| Enteritidis 1171/S/09 | -7.70 | 58.21 | 48.43 | 64.68 | 79.68 | 56.35 |
| Enteritidis 1535/S/09 | -7.70 | 53.28 | 48.43 | 59.20 | 75.90 | 53.68 |
| Enteritidis 476PP2022 | -4.62 | 29.30 | 72.14 | 32.56 | 74.73 | 52.85 |
| Enteritidis 1446/S/09 K31 | -7.01 | 38.88 | 53.73 | 43.20 | 68.72 | 48.60 |
| Enteritidis 212PP2021 | -8.75 | 50.65 | 40.35 | 56.27 | 68.03 | 48.11 |
| Enteritidis 2050/S/09 K4 nr2 | -7.70 | 38.59 | 48.43 | 42.88 | 64.66 | 45.73 |
| Enteritidis 838/S/09 B | -7.70 | 38.28 | 48.43 | 42.53 | 64.42 | 45.56 |
| Typhimurium ATCC 13311 | -7.70 | 35.71 | 48.43 | 39.67 | 62.45 | 44.17 |
| Enteritidis 1445/S/09 K46 | -8.12 | 35.29 | 45.22 | 39.21 | 59.80 | 42.29 |
| Enteritidis 488PP2023 | -7.11 | 11.84 | 53.01 | 13.15 | 47.50 | 33.59 |
| Hadar 817 | -8.12 | 14.09 | 45.22 | 15.66 | 43.57 | 30.82 |
| Typhimurium 1567/S/09 | -8.78 | 16.09 | 40.13 | 17.88 | 41.42 | 29.29 |
| Typhimurium 960/S/09 | -8.12 | 4.13 | 45.22 | 4.59 | 35.95 | 25.42 |
| Paratyphi A ATCC 19150 | -10.19 | 18.29 | 29.29 | 20.33 | 35.24 | 24.92 |
| Brandenburg 584 | -9.78 | 13.71 | 32.44 | 15.23 | 34.01 | 24.05 |
| Enteritidis KK9 | -11.78 | 17.80 | 17.06 | 19.78 | 25.99 | 18.38 |
| Enteritidis KK10 | -12.86 | 22.36 | 8.77 | 24.84 | 23.47 | 16.60 |
| Infantis ZK5 | -10.53 | -0.51 | 26.68 | -0.57 | 18.95 | 13.40 |
| Typhimurium 1081/S/09 K1 | -11.78 | 6.51 | 17.06 | 7.23 | 17.35 | 12.27 |
| Typhimurium 1578/S/09 | -12.86 | 10.84 | 8.77 | 12.04 | 14.65 | 10.36 |
| Enteritidis KK11 | -13.56 | 14.07 | 3.36 | 15.64 | 13.21 | 9.34 |
| Typhimurium 531 | -13.18 | 3.23 | 6.35 | 3.59 | 7.07 | 5.00 |
| Typhimurium 1785/09 | -13.86 | 7.52 | 1.04 | 8.36 | 6.51 | 4.61 |
| Virchow 1188/S/09 K10 | -13.86 | 2.48 | 1.04 | 2.76 | 2.66 | 1.88 |
| Infantis 144 | -13.86 | -0.38 | 1.04 | -0.42 | 0.47 | 0.33 |
| Virchow 1175/S/09 | -13.86 | -2.00 | 1.04 | -2.22 | -0.77 | -0.55 |
| Infantis 1021/S/09 | -13.86 | -2.00 | 1.04 | -2.22 | -0.77 | -0.55 |
| Virchow GAP K2 | -13.86 | -3.74 | 1.04 | -4.16 | -2.11 | -1.49 |

### Representativeness of *Salmonella* strains

All strains were compared in terms of their average nucleotide identity (ANI). The analysis revealed that the collection is diverse in the genomic context. The minimum ANI value between the serovars was found to be 98.22%. Average nucleotide identity parameter, indicated also of the diversity within serovars, with the ANI values ranging from 99.77-100% for 55 Enterica, 99.77-99.98% for Typhimurium, 99.89-99.96% for Infantis and 99.95-99.99% for Virchow serovar strains. The complete ANI heatmap for all 72 *Salmonella enterica* strains used in the study is given in **Figure 3**, whereas for the subset of 55 *Salmonella enterica* ser. Enteritidis strains in **Supplementary Figure S3**.

**Figure 3.**
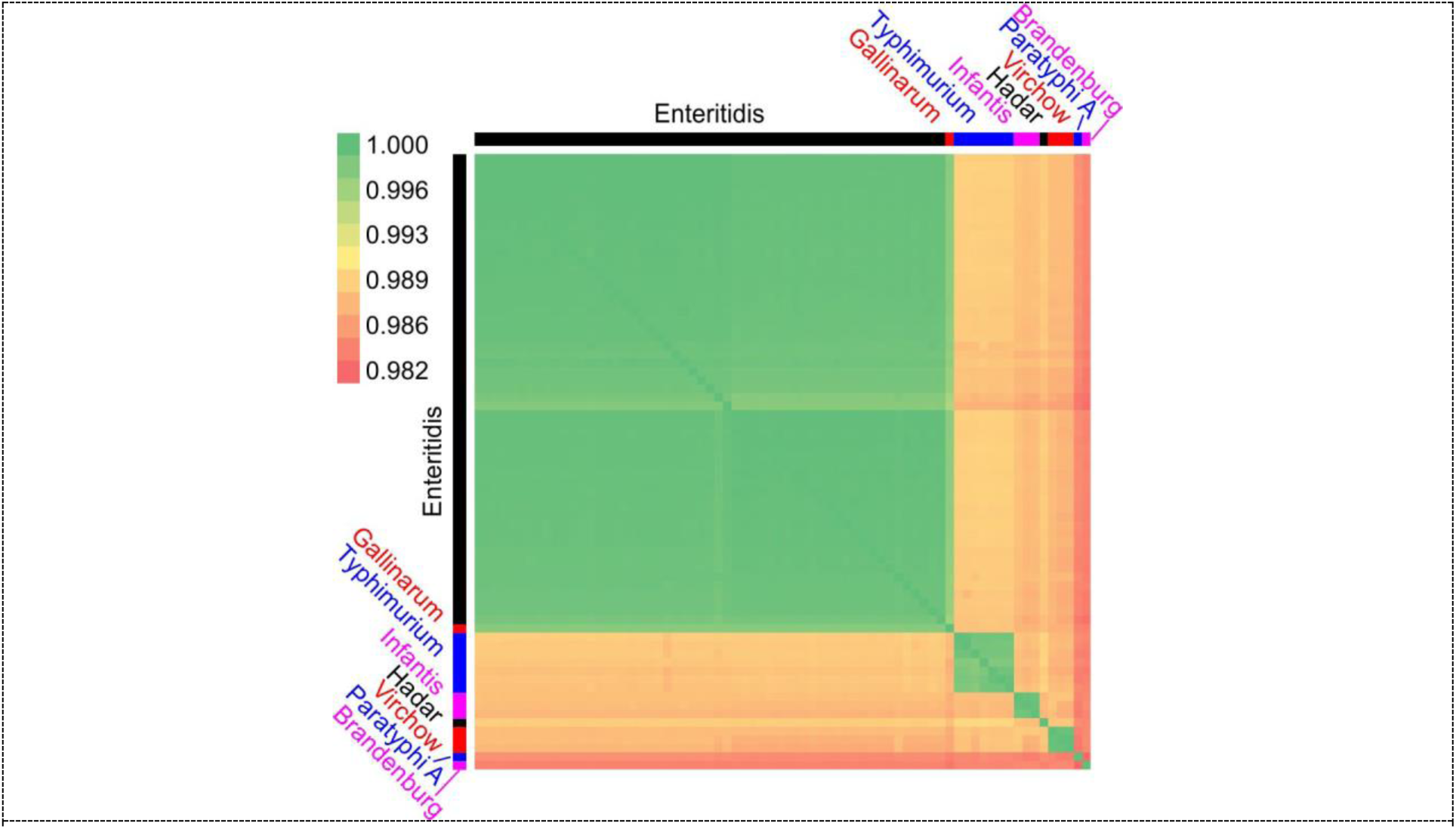
Heatmap of the average nucleotide identity of the genomes of 72 *Salmonella* strains tested in this study.

As expected, bioinformatics analysis revealed that the greatest differences in genomic sequences are observed between different serovars. Further analyses were performed only on *Salmonella* Enteritidis strains as those were the most numerous in this collection. All *S.* Enteritidis strains were proved to be unique, however, to addresses the problem of representativeness of strains employed in the study for expected *Salmonella* strains variability in poultry sites in different geographies, the phylogenetic tree of these strains was created using the Core Genome Multilocus Sequence Typing (cgMST), which was constructed basing on 3002 loci. Resulting tree of the Enteritidis strains is shown in **Figure 4**. Analysis indicated three strains being distinctly different from the other in the dataset - 212PP2021 (originating from Spain), 489PP2023 (Brazil) and 472PP2022 (Poland) and two other strains clustering separately – 093PP2018 from Ukraine and 488PP2023 from Egypt. Also, two isolates from Turkey and one strain from Netherlands are different from the most *Salmonella enterica* ser. Enteritidis strains used in the study, being most similar with Brazilian strain 494PP2023, ATCC 13076 strain and one of the Polish strains 249. Although strains from Poland are major collection component (76%) the distance between some of strains isolated in Poland is in many instances greater than the distance between strains from Poland and strains from other geographic locations.

**Figure 4.**
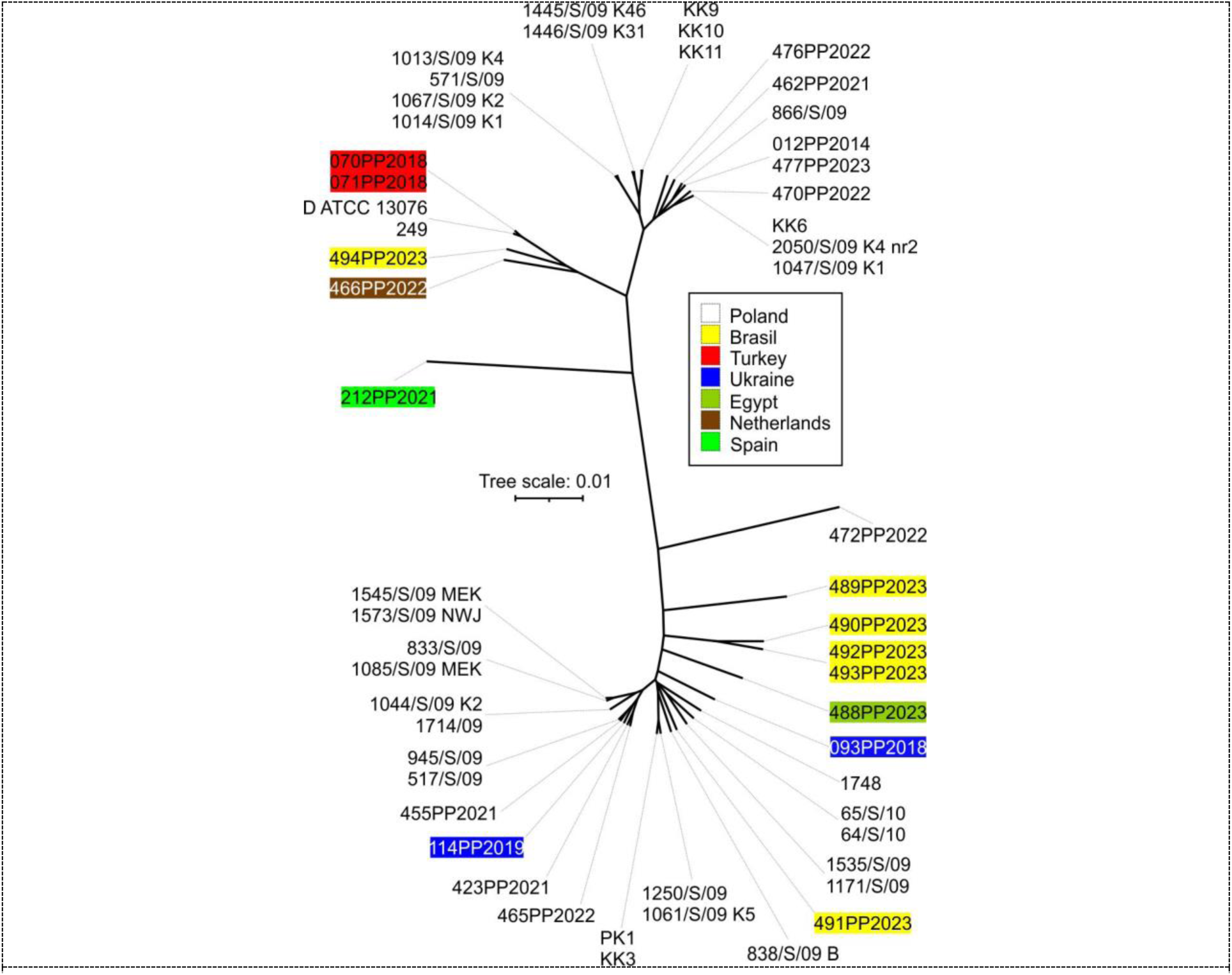
Phylogenetic tree for 55 *Salmonella enterica* ser. Enteritidis strains used in the study, constructed using cgMLST profiles. Strains obtained from outside of Poland are labeled with country name and color highlight. Based on backbone created with iTol v6 tool.

Additionally, putative antibiotic resistance genotypes were detected by AMRFinderPlus tool in tested strains. Two multidrug efflux pump genes were identified - ***mds*A** and ***mds*B**, and for 32 of them these two genes were the only AMR genes identified. Remaining Enteritidis strains revealed the presence of 3 or 4 AMR genes consisting of: β-lactam antibiotic resistance genes - ***bla*TEM, *bla*TEM-1**, tetracycline resistance - ***tet*(A)**. There were also four different point mutations (D87N, D87Y, S83F, S83Y) found in DNA gyrase subunit A **(*gyr*A)** gene, which correspond to quinolone (and/or triclosan) resistance, as well as unique (present for only one Egyptian *Salmonella enterica* ser. Enteritidis 488PP2023 strain) point mutation (R717L) within multidrug efflux RND transporter permease subunit ***acr*B** gene conferring resistance to macrolide antibiotic azithromycin.

For virulence factors profiles identification VFDB tool was used. For fifty-five *Salmonella* Enteritidis strains from 165 to 325 VF per strain were identified, with the median of 174 virulence factors per strain. Majority of the strains of this serovar (n=24) had 175 virulence factors identified which mainly consisted of genes regarding fimbrial and nonfimbrial adherence determinants and secretion systems. In contrast, a single strain 476PP2022, for which 325 virulence factors were identified was checked for possible contamination, but none was determined (over 98% of sequencing reads assigned to *Salmonella* genus, no 16S RNA fragments different from *Salmonella*). Overview of the AMR genes and virulence factors in Enteritidis strains is presented in **Table 6**.

**Table 6.** AMR genes and virulence factors in tested *S.* Enteritidis strains

| AMR genes |  |  |  | AMR genes |  |  |  |
| --- | --- | --- | --- | --- | --- | --- | --- |
| Strain | n | names | n of virulence factors identified | Strain | n | names | n of virulence factors identified |
| Enteritidis 012PP2014 | 2 | *mdsA, mdsB* | 174 | Enteritidis 466PP2022 | 2 | *mdsA, mdsB* | 174 |
| Enteritidis 070PP2018 | 2 | *mdsA, mdsB* | 174 | Enteritidis 470PP2022 | 2 | *mdsA, mdsB* | 174 |
| Enteritidis 071PP2018 | 2 | *mdsA, mdsB* | 174 | Enteritidis 472PP2022 | 2 | *mdsA, mdsB* | 166 |
| Enteritidis 093PP2018 | 3 | *gyrA_S83F, mdsA, mdsB* | 175 | Enteritidis 476PP2022 | 2 | *mdsA, mdsB* | 325 |
| Enteritidis 1013/S/09 K4 | 2 | *mdsA, mdsB* | 174 | Enteritidis 477PP2023 | 2 | *mdsA, mdsB* | 174 |
| Enteritidis 1014/S/09 K1 | 2 | *mdsA, mdsB* | 174 | Enteritidis 488PP2023 | 3 | *acrB_R717L, mdsA, mdsB* | 174 |
| Enteritidis 1044/S/09 K2 | 3 | *gyrA_S83F, mdsA, mdsB* | 175 | Enteritidis 489PP2023 | 2 | *mdsA, mdsB* | 174 |
| Enteritidis 1047/S/09 K1 | 2 | *mdsA, mdsB* | 174 | Enteritidis 490PP2023 | 3 | *gyrA_D87N, mdsA, mdsB* | 175 |
| Enteritidis 1061/S/09 K5 | 3 | *gyrA_D87Y, mdsA, mdsB* | 175 | Enteritidis 491PP2023 | 2 | *mdsA, mdsB* | 175 |
| Enteritidis 1067/S/09 K2 | 2 | *mdsA, mdsB* | 174 | Enteritidis 492PP2023 | 3 | *gyrA_D87N, mdsA, mdsB* | 174 |
| Enteritidis 1085/S/09 MEK | 3 | *gyrA_S83Y, mdsA, mdsB* | 175 | Enteritidis 493PP2023 | 3 | *gyrA_D87N, mdsA, mdsB* | 175 |
| Enteritidis 114PP2019 | 3 | *gyrA_S83Y, mdsA, mdsB* | 175 | Enteritidis 494PP2023 | 2 | *mdsA, mdsB* | 165 |
| Enteritidis 1171/S/09 | 3 | *mdsA, mdsB, tet(A)* | 178 | Enteritidis 517/S/09 | 3 | *gyrA_S83Y, mdsA, mdsB* | 175 |
| Enteritidis 1250/S/09 | 4 | *blaTEM-1, gyrA_D87Y, mdsA, mdsB* | 175 | Enteritidis 571/S/09 | 2 | *mdsA, mdsB* | 174 |
| Enteritidis 1445/S/09 K46 | 2 | *mdsA, mdsB* | 174 | Enteritidis 64/S/10 | 2 | *mdsA, mdsB* | 175 |
| Enteritidis 1446/S/09 K31 | 2 | *mdsA, mdsB* | 173 | Enteritidis 65/S/10 | 2 | *mdsA, mdsB* | 175 |
| Enteritidis 1535/S/09 | 3 | *mdsA, mdsB, tet(A)* | 178 | Enteritidis 833/S/09 | 3 | *gyrA_S83Y, mdsA, mdsB* | 175 |
| Enteritidis 1545/S/09 MEK | 3 | *gyrA_S83Y, mdsA, mdsB* | 175 | Enteritidis 838/S/09 B | 2 | *mdsA, mdsB* | 173 |
| Enteritidis 1573/S/09 NWJ | 3 | *gyrA_S83Y, mdsA, mdsB* | 175 | Enteritidis 866/S/09 | 2 | *mdsA, mdsB* | 174 |
| Enteritidis 1714/09 | 3 | *blaTEM, mdsA, mdsB* | 175 | Enteritidis 945/S/09 | 3 | *gyrA_S83Y, mdsA, mdsB* | 175 |
| Enteritidis 1748 | 2 | *mdsA, mdsB* | 175 | Enteritidis D ATCC 13076 | 2 | *mdsA, mdsB* | 174 |
| Enteritidis 2050/S/09 K4 nr2 | 2 | *mdsA, mdsB* | 173 | Enteritidis KK10 | 2 | *mdsA, mdsB* | 174 |
| Enteritidis 212PP2021 | 2 | *mdsA, mdsB* | 175 | Enteritidis KK11 | 2 | *mdsA, mdsB* | 174 |
| Enteritidis 249 | 2 | *mdsA, mdsB* | 174 | Enteritidis KK3 | 3 | *gyrA_D87Y, mdsA, mdsB* | 175 |
| Enteritidis 423PP2021 | 3 | *gyrA_S83Y, mdsA, mdsB* | 175 | Enteritidis KK6 | 2 | *mdsA, mdsB* | 174 |
| Enteritidis 455PP2021 | 3 | *gyrA_S83Y, mdsA, mdsB* | 175 | Enteritidis KK9 | 2 | *mdsA, mdsB* | 174 |
| Enteritidis 462PP2021 | 2 | *mdsA, mdsB* | 174 | Enteritidis PK1 | 3 | *gyrA_D87Y, mdsA, mdsB* | 175 |
| Enteritidis 465PP2022 | 3 | *gyrA_S83Y, mdsA, mdsB* | 175 |  |  |  |  |

To verify *in silico* observations, the collection of Enteritidis strains was tested for susceptibility to Sulphamethoxazole/Trimethoprim, Enrofloxacin, Gentamycin and Ampicillin. None of the strains tested showed resistance to Sulphamethoxazole/Trimethoprim. 4 strains showed partial resistance to Enrofloxacin (susceptible at elevated concentrations), 6 strains showed resistance to Gentamycin and 2 strains showed resistance to Ampicillin. Only 2 strains showed resistance or partial resistance to two types of antibiotics. The results (zone diameter of inhibition) of antibiotic-resistant strains are presented in **Table 7**. The results for susceptible strains and for Sulphamethoxazole/Trimethoprim are omitted, the full results are presented in the **Supplementary Table S2**.

**Table 7.** The results of the antibiotic resistance of tested *S.* Enteritidis strains. R – resistant. ATU – susceptible, increased exposure.

| Strain | Enrofloxacin<br>(ENR 5) | Gentamycin<br>(CN 10) | Ampicillin<br>(AMP 10) |
| --- | --- | --- | --- |
| Enteritidis 114PP2019 | 18 (ATU) | 16 (R) |  |
| Enteritidis 1250/S/09 |  |  | 0 (R) |
| Enteritidis 1445/S/09 K46 |  | 16 (R) |  |
| Enteritidis 1545/S/09 MEK | 22 (ATU) |  |  |
| Enteritidis 455PP2021 | 20 (ATU) |  |  |
| Enteritidis 490PP2023 |  | 16 (R) |  |
| Enteritidis 517/S/09 |  | 14 (R) | 12 (R) |
| Enteritidis 571/S/09 |  | 16 (R) |  |
| Enteritidis 833/S/09 | 22 (ATU) |  |  |
| Enteritidis KK9 |  | 16 (R) |  |

### The correlation of susceptibility results with AMR and virulence profiles of *Salmonella* Enteritidis strains

Results of in-depth analysis of susceptibility of Enteritidis strains to BAFASAL^®^ is presented in **Figure 5**. Histogram of normalized CLS values presented in **Figure 4A** shows that over 65% of Enteritidis strains have reached normalized CLS higher than 75% and only 3 strains fall below 30%. Two of the tested strains, marked in **Figure 5**, were previously used in the *in vivo* BAFASAL^®^ efficacy experiments [33, 45]. Subsequently, Enteritidis strains were divided into two groups based on antibiotic resistance tested *in vitro* (both resistant and susceptible at increased exposure strains were treated as positive) or number of antibiotic resistance genes or number of virulence genes (found *in silico* from sequencing data). Mann-Whitney test was performed to check whether median of the CLS values in groups significantly differs (**Figure 5B**). No significant differences were found suggesting strongly that susceptibility towards BAFASAL^®^ is not correlated with either antibiotic resistance or virulence of the *Salmonella* strain.

**Figure 5.**
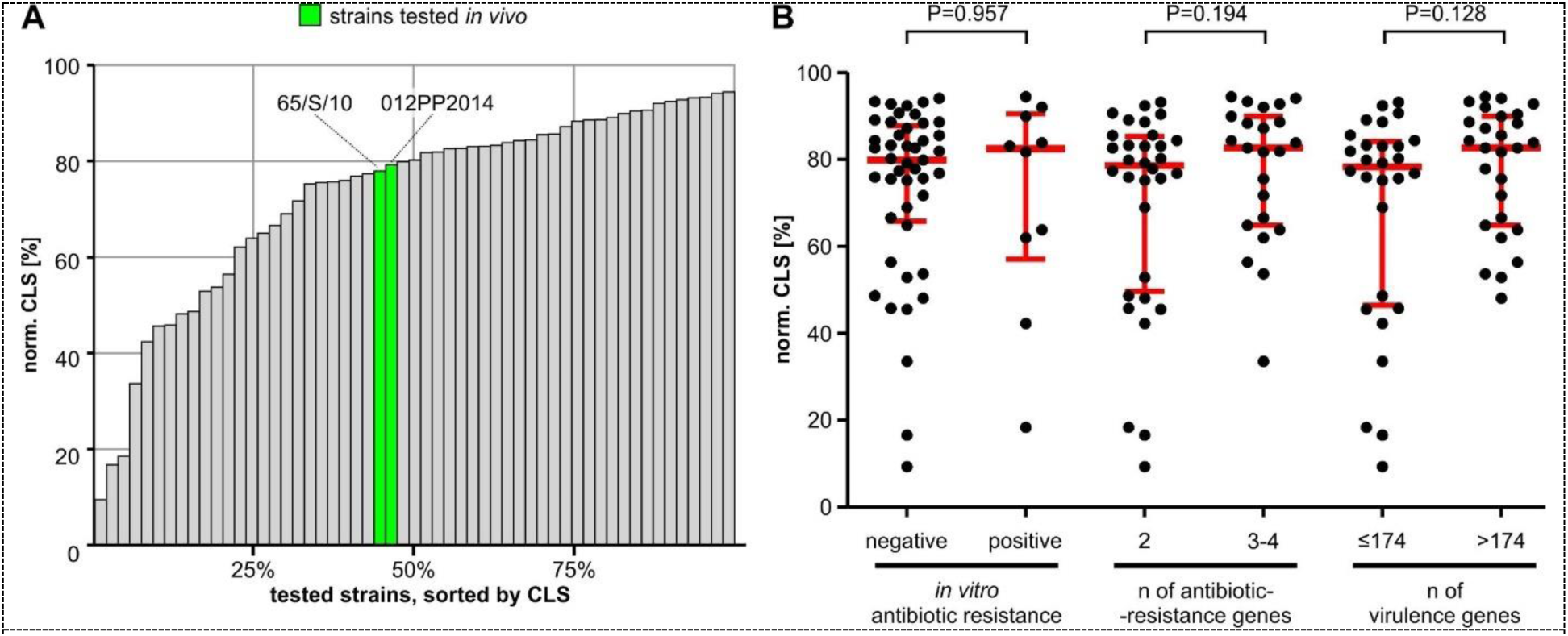
CLS analysis for Enteritidis strains. **A:** histogram of the normalized CLS values, sorted in ascending order; **B:** Comparison of CLS values in groups, depending on *in vitro* antibiotic resistance or number of antibiotic-resistance genes or virulence genes.

## DISCUSSION

During this study a collection of 72 unique *Salmonella* strains was sequenced, analyzed bioinformatically and tested *in vitro* for susceptibility to phage-mediated lysis with BAFASAL^®^ preparation. In addition, susceptibility of all *Salmonella* strains to panel of antibiotics was also experimentally determined. This bacterial collection consisted of 8 different serovars of *Salmonella enterica* with 55 strains representing serovar Enteritidis. This significant overrepresentation of Enteritidis was intentional as intended BAFASAL^®^ application is control of *Salmonella* Enteritidis in poultry.

Genomic analysis confirmed that all bacterial strains are unique. The geographic origin of Enteritidis strains in collection is heterogenous although strains isolated in Poland are overrepresented. However, considering level of genetic similarity between Enteritidis strains from Poland and strains from distant geographic location (**Figure 4**) Enteritidis part of bacterial collection seems to well represent global diversity of this serovar in poultry. All bacterial strains were also analyzed for genetic variability (**Figure 3**), antibiotic resistance, and presence of pathogenicity mediating genes (**Table 6**).

To further validate bioinformatic predictions, *in vitro* antibiotic sensitivity assays were performed (**Table 7**). Panel of antibiotics consisting of Sulphamethoxazole/Trimethoprim, Enrofloxacin, Gentamycin and Ampicillin have been used. Sulphamethoxazole/Trimethoprim was chosen on the basis of EUCAST recommendations for *Salmonella* testing, Enrofloxacin as the most popular in poultry production, while Gentamycin and Ampicillin as beta-lactams and aminoglycosides representatives, respectively in accordance outcome of *in silico* analysis of antibiotic resistance genes in genomes of tested strains. In general, *in vitro* results does not correlate strictly with identified AMR genes. In many cases drug-specific AMR genes present in the genomes does not confer to the resistance *in vitro.* On the other hand, the most abundant AMR genes in tested strains are multidrug resistance pumps coded by *mds*A, *mds*B, and these genes seems to be also the most impactful. None of Enteritidis strains carries a gene of resistance of sulfonamide and no corresponding resistance towards Sulphamethoxazole/Trimethoprim was observed *in vitro*. Only 4 strains showed moderate resistance to Enrofloxacin and all 4 of them carries the *gyr*A_S83Y mutation, which might suggest that expression of this gene variant is involved in the resistance mechanism. However, 6 remaining strains sharing the same mutation have not shown resistance. None of Enteritidis strains expressed genes corresponding to specific resistance of aminoglycosides. Thus observed resistance of 6 strains to Gentamycin, was most likely caused by multidrug resistance efflux pumps coded by *mds*A, *mds*B genes. Lastly, 2 strains showed resistance of Ampicillin, one of which was carrying beta-lactamase gene *bla*TEM-1 and the other had no drug-specific gene. In the latter case, probably efflux pumps were causing observed AMR, especially that in this case in higher concentrations antibiotic action was clearly visible (radius of inhibition zone of 12 mm) and beta-lactamase-positive strain was 100% resistant (0 mm of inhibition). The only other Enteritidis strain carrying beta-lactamase gene (*bla*TEM) showed no resistance *in vitro*. Our observations are similar to previously published studies [46, 47, 48], where *in vitro* AMR resistance is only partially correlated with the presence of AMR genes because of the presence of silent AMR genes that did not confer actual resistance and expression of efflux pumps being important factor of multidrug resistance phenotype.

Host range of BAFASAL^®^ preparation was determined by testing susceptibility of all *Salmonella* strains using combination of serial dilution spot test and spectrophotometric analysis of bacterial growth kinetics. These two methods are broadly accepted as appropriate tools for determination of host range of bacteriophages and bacteriophage cocktails [14, 19, 20, 25]. These methods are also recognized as appropriate for such studies by EMA in the guideline for veterinary medicinal products for bacteriophage therapy [49]. It is important to acknowledge that these methods measure bacterial response in different type of culture (semisolid and liquid) and using different experimental endpoints. Additionally, both assays differ in their capabilities to effectively quantify the tested strains susceptibility. In the case of the spot test, multiple phage concentrations are tested but many strains achieve the same result as number of possible outcomes is limited, also no information about the kinetics of undergoing lysis is collected. In contrast, the spectrophotometric assay allows to compare the kinetics, so slight differences in the growth rate of the culture could be observed but is typically limited to one concentration of the phages that may be too low or too high to show differences between susceptibilities of different bacterial strains.

Results from two *in vitro* assays were generally comparable (**Table 3** and **Table 4**). 86.1% of tested strains achieved the same result in both tests. This is similar to previously published studies employing spot assay and spectrophotometric assay in parallel. Xie et al. observed close agreement in 74% cases testing 15 bacteriophage strains against 20 strains of *Salmonella* [20]. Similar observation was made by Cooper et al. with 4 bacteriophage strains and 14 Pseudomonas aeruginosa strains. In both reports authors indicated correlation between results obtained with each method but they did not performed further mathematical analysis of this correlation [24].

One limitation in comparison of results of these assays was difference in measurement scales with spectrophotometric assay using continuous scale, and spot test series of nominal observations (no effect or different types of lysis). To overcome this problem, we transformed raw results from the spot test using the same statistical approach which is used in counting microorganisms in the serial dilutions method i.e. the most probable number method (MPN). By this method one is able to calculate the theoretical titer of the microorganisms in a solution. In our case, because titer of phages used was already known and equal among all the experiments, we assumed that outcome of this estimation is corresponding to susceptibility of the tested strain towards tested phage preparation. To conveniently compare different strains/phage preparations one can divide this estimated theoretical number of phages by the known titer used. We named this metric the Effective Concentration Threshold (ECT). The lesser the ECT value the more susceptible bacteria is towards the tested phage. Another problem in transforming outcome of spot assay into numerical values were different types of lysis observed in spot test suggesting possibility of extracting more information that simple present/absent observation. To explore further this concept two analytical components were created, one based on any lytic event and the other on clear lysis (the strongest lytic effect observed) only. Subsequently, to confirm whether this approach is correct and to test what should be relative weight of this two components in the final result, computer optimization against correlation coefficient between spot test and spectrophotometric test results was performed (**Figure 1**). The premise to optimize the spot assay score equation this way was built on assumption that in fact, results from both tests are mainly based on the strains susceptibility which we were trying to measure. Thus, with large enough dataset this approach should produce a valuable equation. Finally, complete equation for the spot test score consists of two ECT components (one for any lysis and one for clear lysis only), both logarithmized (to linearize the data from several orders of magnitude) and added. Lastly, the minus sign was added just to keep the direct proportional relation with the spectrophotometric test results.

Then, when we finally compared the correlation of the results of both tests, we obtained an R^2^ value of 82.4%. Subsequently, using a mathematical approach similar to the principal component analysis method, i.e., by calculating the scalar product of the coordinates of each point and a unit vector parallel to the regression line, we sorted the strains in ascending order of sensitivity to bacteriophage lysis *in vitro*. We believe that this new, holistic metric reflects the influence of both the bacteriophage concentration threshold component (spot assay) and the component of the intensity of lysis or growth rate limitation (spectrophotometric assay). The method we presented is suitable for quantitative comparison of bacterial strain sensitivity to phage lysis *in vitro*. By using the calibration values, we obtained from screening our strain collection, and the equations we propose, one should be able to compare small groups of strains or even to perform pairwise comparisons, without the need of further calibration. However, we believe that the greatest potential of this method lies in sorting and quantifying data from large screening studies. In this case, we recommend that researchers perform calibration on their own dataset (as it is descibed in the paragraph “Analysis of the correlation between two *in vitro* tests” in the results section). It is also worth mentioning that while in the case of the spot test, small differences (± order of magnitude) in the initial bacteriophage concentration should not affect the obtained results, in the case of the spectrophotometric test this is a factor that should be taken into account when comparing the results from several studies, as this effect cannot be easily predicted or normalized.

Determination of the specificity of BAFASAL^®^ preparation using both *in vitro* methods generated consistent results with significant correlation between methods. Assessment of host range using new metric of Combined Lytic Score (CLS) showed that 50% of the tested strains have reached normalized CLS of over 75% (**Table 5**, **Figure 2**). When focusing only on Enteritidis strains, over 67% of Enteritidis strains have reached normalized CLS higher than 75% and only 4 strains fall below 40%. Importantly, susceptibility of Enteritidis strains to BAFASAL^®^ preparation does not depend on strain pathogenicity and antibiotic resistance (**Figure 5**). Considering CLS observed for the validated *in vivo* strain Enteritidis 012PP2014 [45] expected host range *in vivo* is at least at the level of 55% (30 out of 55 strains). This is based on very conservative assumption that only strains with CLS equal or higher to Enteritidis 012PP2014 will be susceptible *in vivo*. Using less stringent condition that CLS value of 70% is enough to consider strain susceptibility, the host range *in vivo* would be predicted at the level of 69% (38 out of 55 strains). It is worth to underline that the data and methodology presented in this study were positively assessed by EFSA and led to the decision that BAFASAL® has the potential to reduce environmental contamination with *S.* Enteritidis when used in feed and water for all poultry species [50].

## Supplementary file 1

**Table S1.**
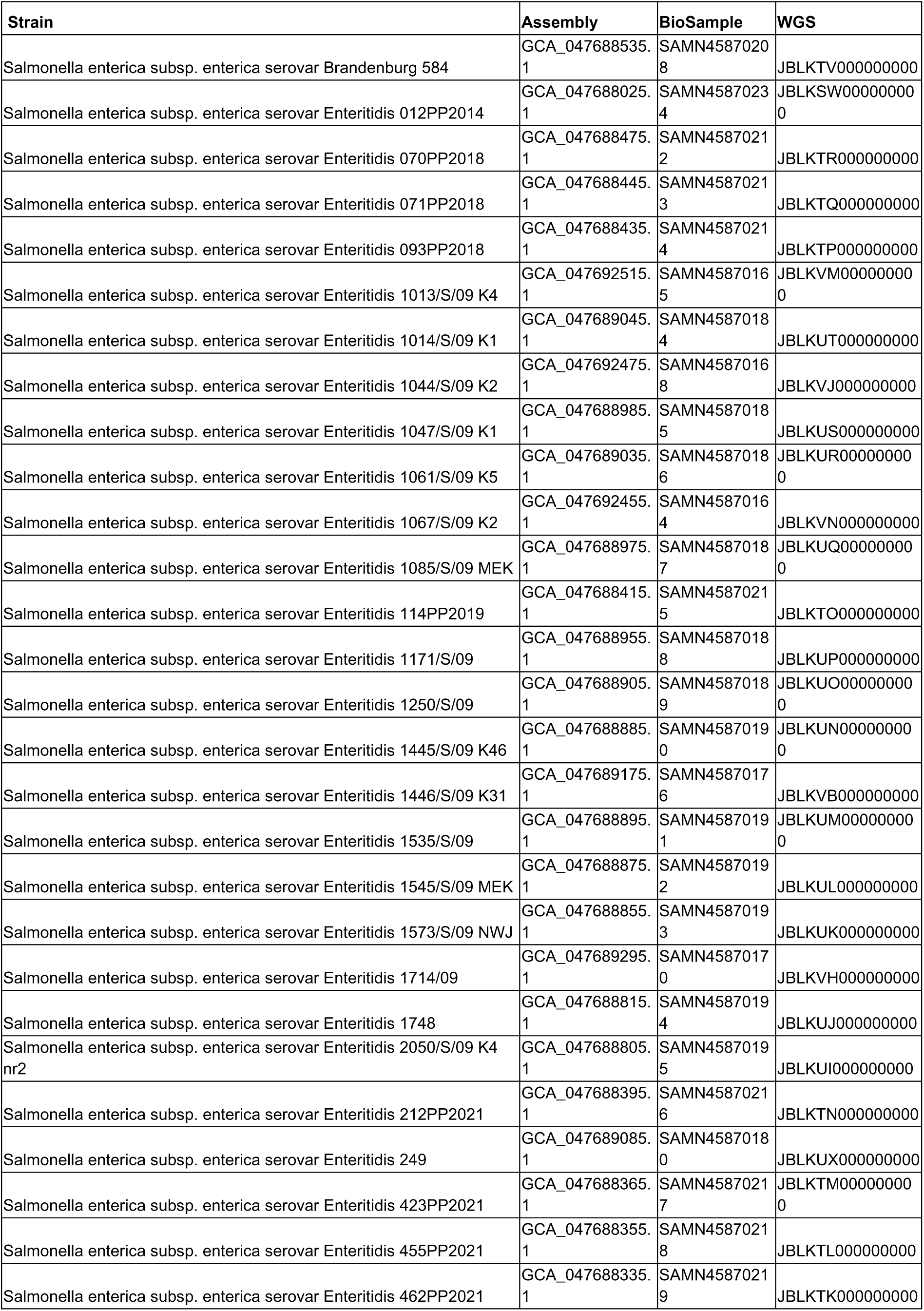
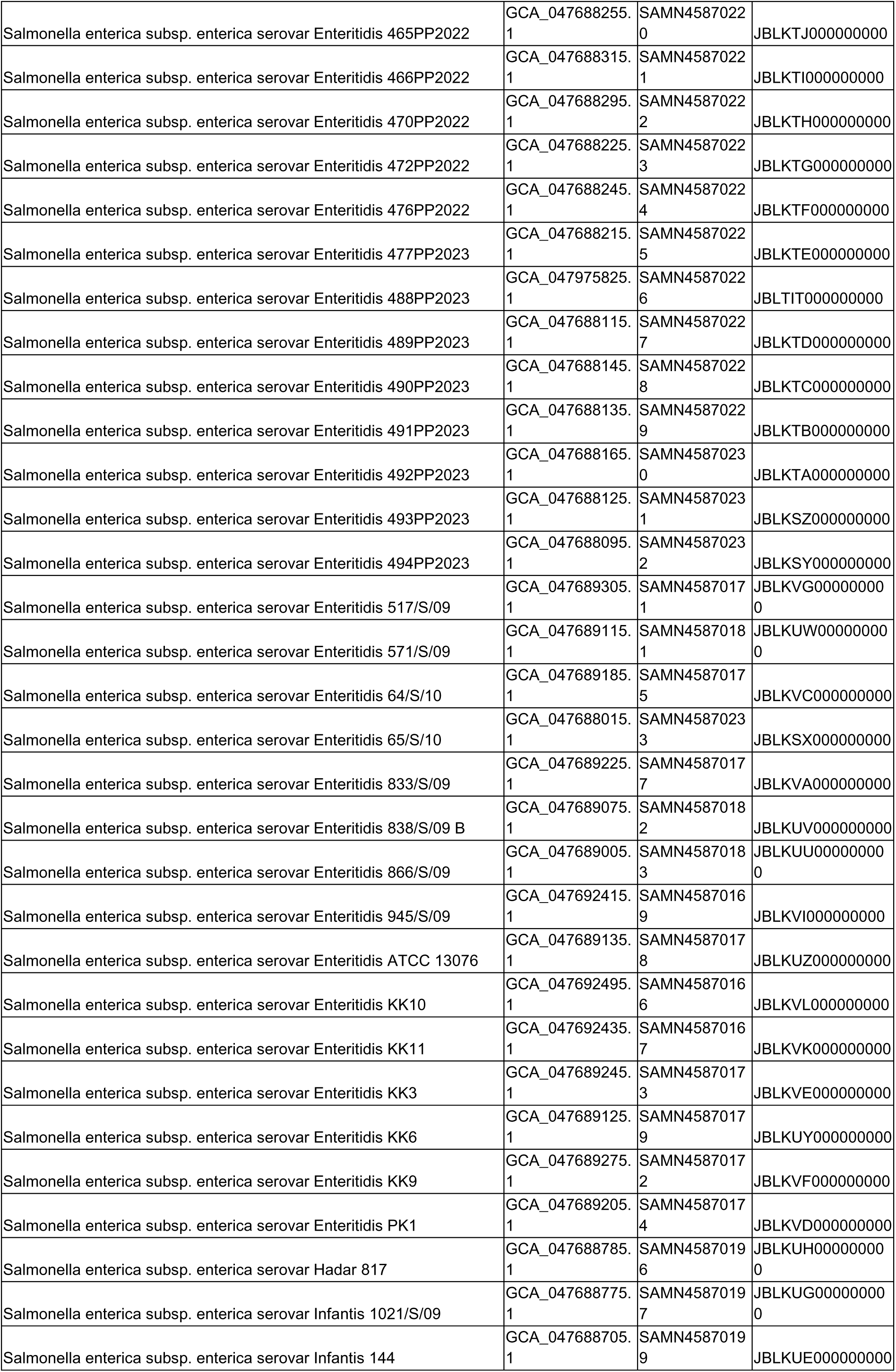
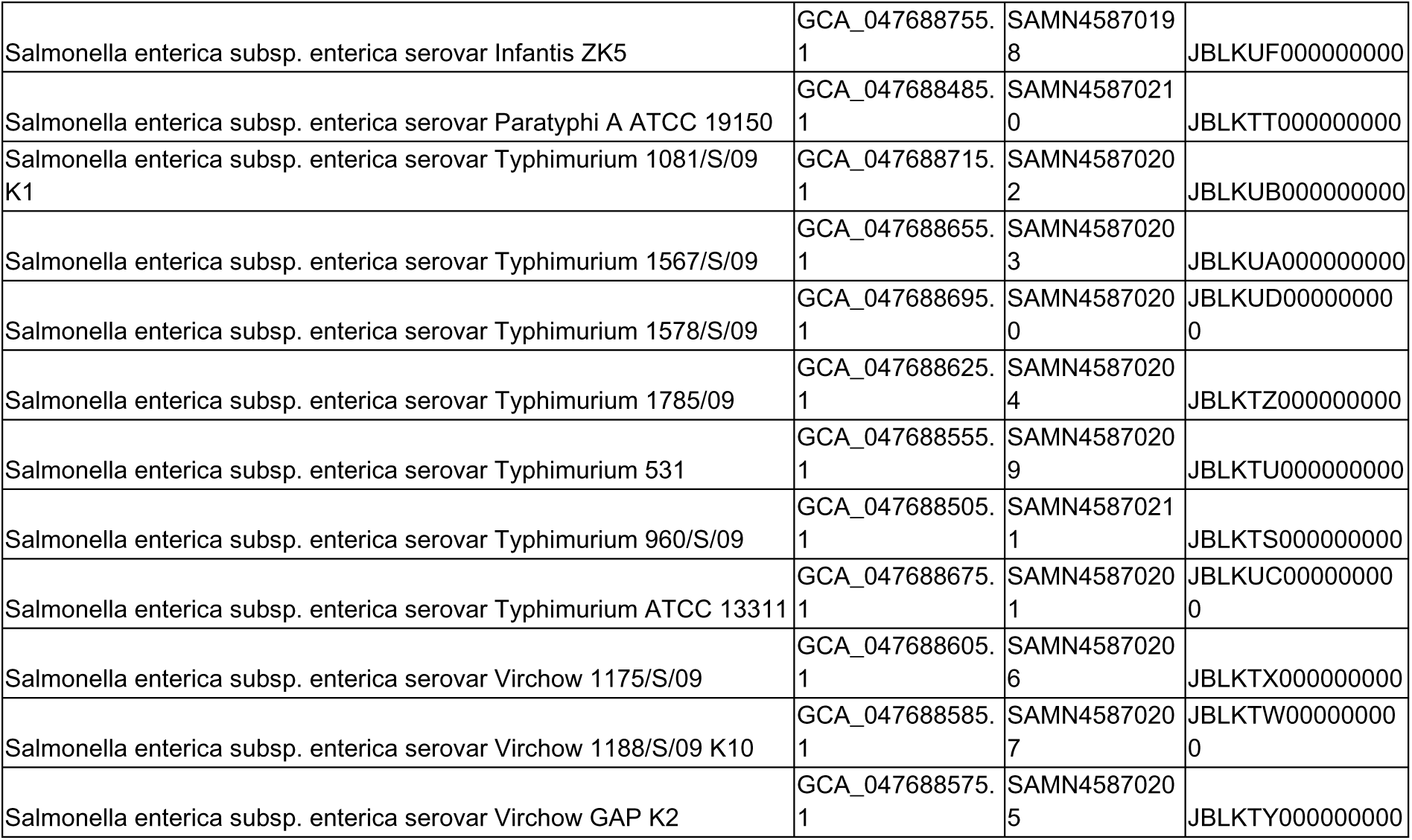
Accession numbers for the genomic sequences of the used *Salmonella* strains

## Supplementary file 2

**The full results of the *in vitro* AMR tests**

**Table S2.** The results of the antibiotic resistance of tested *S. enterica* strains

| *Salmonella enterica* strain | Bacterial growth inhibition zone diameter [mm] |  |  |  |
| --- | --- | --- | --- | --- |
|  | Sulphamethoxazole/<br>Trimethoprim (SXT 25)<br>S ≥ 14 mm<br>R < 11 mm<br>ATU 12-13 mm | Enrofloxacin (ENR 5)<br>S ≥ 23 mm<br>R < 16 mm<br>ATU 17-22 mm | Gentamycin (CN 10)<br>S ≥ 17 mm<br>R < 17 mm | Ampicillin (AMP 10)<br>S ≥ 14 mm<br>R < 14 mm |
| Gallinarum 001PP2015 | 35 (S) | 30 (S) | 22 (S) | 33 (S) |
| Enteritidis 1067/S/09 K2 | 32 (S) | 34 (S) | 20 (S) | 24 (S) |
| Enteritidis 1013/S/09 K4 | 34 (S) | 34 (S) | 20 (S) | 24 (S) |
| Enteritidis KK10 | 32 (S) | 32 (S) | 20 (S) | 24 (S) |
| Enteritidis KK11 | 30 (S) | 32 (S) | 20 (S) | 24 (S) |
| Enteritidis 1044/S/09 K2 | 32 (S) | 30 (S) | 20 (S) | 28 (S) |
| Enteritidis 945/S/09 | 30 (S) | 24 (S) | 20 (S) | 28 (S) |
| Enteritidis 1714/09 | 28 (S) | 32 (S) | 20 (S) | 26 (S) |
|  | Sulphamethoxazole/<br>Trimethoprim (SXT 25)<br>S≥14 mm<br>R<11 mm<br>ATU 12-13 mm | Enrofloxacin (ENR 5)<br>S≥23 mm<br>R<16 mm<br>ATU 17-22 mm | Gentamycin (CN 10)<br>S≥17 mm<br>R<17 mm | Ampicillin (AMP 10)<br>S≥14 mm<br>R<14 mm |
| Enteritidis 517/S/09 | 32 (S) | 26 (S) | 14 (R) | 12 (R) |
| Enteritidis KK9 | 30 (S) | 32 (S) | 16 (R) | 24 (S) |
| Enteritidis KK3 | 28 (S) | 26 (S) | 20 (S) | 26 (S) |
| Enteritidis PK1 | 28 (S) | 26 (S) | 18 (S) | 24 (S) |
| Enteritidis 64/S/10 | 30 (S) | 30 (S) | 18 (S) | 26 (S) |
| Enteritidis 1446/S/09 K31 | 30 (S) | 26 (S) | 18 (S) | 24 (S) |
| Enteritidis 833/S/09 | 30 (S) | 22 (ATU) | 18 (S) | 24 (S) |
| Enteritidis D ATCC 13076 | 30 (S) | 30 (S) | 18 (S) | 26 (S) |
| Enteritidis KK6 | 30 (S) | 30 (S) | 20 (S) | 24 (S) |
| Enteritidis 249 | 30 (S) | 34 (S) | 18 (S) | 24 (S) |
| Enteritidis 571/S/09 | 30 (S) | 30 (S) | 16 (R) | 26 (S) |
| Enteritidis 838/S/09 B | 30 (S) | 30 (S) | 20 (S) | 22 (S) |
| Enteritidis 866/S/09 | 30 (S) | 28 (S) | 20 (S) | 24 (S) |
| Enteritidis 1014/S/09 K1 | 32 (S) | 32 (S) | 18 (S) | 26 (S) |
| Enteritidis 1047/S/09 K1 | 30 (S) | 30 (S) | 18 (S) | 26 (S) |
| Enteritidis 1061/S/09 K5 | 32 (S) | 24 (S) | 18 (S) | 28 (S) |
| Enteritidis 1085/S/09 MEK | 32 (S) | 26 (S) | 18 (S) | 28 (S) |
| Enteritidis 1171/S/09 | 30 (S) | 32 (S) | 20 (S) | 24 (S) |
| Enteritidis 1250/S/09 | 32 (S) | 26 (S) | 20 (S) | 0 (R) |
| Enteritidis 1445/S/09 K46 | 32 (S) | 32 (S) | 16 (R) | 26 (S) |
| Enteritidis 1535/S/09 | 30 (S) | 32 (S) | 20 (S) | 28 (S) |
| Enteritidis 1545/S/09 MEK | 30 (S) | 22 (ATU) | 18 (S) | 22 (S) |
| Enteritidis 1573/S/09 NWJ | 34 (S) | 26 (S) | 18 (S) | 22 (S) |
| Enteritidis 1748 | 32 (S) | 32 (S) | 20 (S) | 28 (S) |
| Enteritidis 2050/S/09 K4 nr2 | 30 (S) | 34 (S) | 20 (S) | 30 (S) |
| Hadar 817 | 32 (S) | 26 (S) | 18 (S) | 0 (R) |
| Infantis 1021/S/09 | 28 (S) | 32 (S) | 18 (S) | 0 (R) |
| Infantis Zdul K5 | 30 (S) | 32 (S) | 20 (S) | 30 (S) |
| Infantis 144 | 32 (S) | 32 (S) | 18 (S) | 26 (S) |
| Typhimurium 1578/S/09 | 26 (S) | 24 (S) | 20 (S) | 0 (R) |
| Typhimurium ATCC 13311 | 34 (S) | 34 (S) | 20 (S) | 30 (S) |
| Typhimurium 1081/S/09 K1 | 28 (S) | 24 (S) | 18 (S) | 26 (S) |
| Typhimurium 1567/S/09 | 34 (S) | 32 (S) | 18 (S) | 28 (S) |
| Typhimurium 1785/09 | 32 (S) | 34 (S) | 16 (R) | 26 (S) |
|  | Sulphamethoxazole/<br>Trimethoprim (SXT 25)<br>S≥14 mm<br>R<11 mm<br>ATU 12-13 mm | Enrofloxacin (ENR 5)<br>S≥23 mm<br>R<16 mm<br>ATU 17-22 mm | Gentamycin (CN 10)<br>S≥17 mm<br>R<17 mm | Ampicillin (AMP 10)<br>S≥14 mm<br>R<14 mm |
| Virchow GAP K2 | 30 (S) | 26 (S) | 18 (S) | 26 (S) |
| Virchow 1175/S/09 | 28 (S) | 24 (S) | 18 (S) | 26 (S) |
| Virchow 1188/S/09 K10 | 26 (S) | 20 (ATU) | 14 (R) | 24 (S) |
| Brandenburg 584 | 28 (S) | 30 (S) | 16 (R) | 24 (S) |
| Typhimurium 531 | 26 (S) | 30 (S) | 16 (R) | 0 (R) |
| Paratyphi A ATCC 19150 | 28 (S) | 34 (S) | 20 (S) | 30 (S) |
| Typhimurium 960/S/09 | 28 (S) | 30 (S) | 18 (S) | 22 (S) |
| Enteritidis 070PP2018 | 28 (S) | 30 (S) | 18 (S) | 24 (S) |
| Enteritidis 071PP2018 | 30 (S) | 32 (S) | 18 (S) | 26 (S) |
| Enteritidis 093PP2018 | 30 (S) | 24 (S) | 18 (S) | 26 (S) |
| Enteritidis 114PP2019 | 30 (S) | 18 (ATU) | 16 (R) | 22 (S) |
| Enteritidis 212PP2021 | 30 (S) | 32 (S) | 18 (S) | 22 (S) |
| Enteritidis 423PP2021 | 30 (S) | 24 (S) | 20 (S) | 24 (S) |
| Enteritidis 455PP2021 | 28 (S) | 20 (ATU) | 18 (S) | 24 (S) |
| Enteritidis 462PP2021 | 28 (S) | 30 (S) | 18 (S) | 24 (S) |
| Enteritidis 465PP2022 | 30 (S) | 24 (S) | 20 (S) | 22 (S) |
| Enteritidis 466PP2022 | 30 (S) | 32 (S) | 18 (S) | 26 (S) |
| Enteritidis 470PP2022 | 30 (S) | 30 (S) | 18 (S) | 22 (S) |
| Enteritidis 472PP2022 | 34 (S) | 30 (S) | 22 (S) | 26 (S) |
| Enteritidis 476PP2022 | 30 (S) | 28 (S) | 18 (S) | 26 (S) |
| Enteritidis 477PP2023 | 32 (S) | 30 (S) | 20 (S) | 28 (S) |
| Enteritidis 488PP2023 | 26 (S) | 26 (S) | 18 (S) | 24 (S) |
| Enteritidis 489PP2023 | 28 (S) | 30 (S) | 18 (S) | 26 (S) |
| Enteritidis 490PP2023 | 32 (S) | 26 (S) | 16 (R) | 26 (S) |
| Enteritidis 491PP2023 | 30 (S) | 30 (S) | 18 (S) | 26 (S) |
| Enteritidis 492PP2023 | 30 (S) | 28 (S) | 18 (S) | 28 (S) |
| Enteritidis 493PP2023 | 30 (S) | 26 (S) | 18 (S) | 26 (S) |
| Enteritidis 494PP2023 | 32 (S) | 30 (S) | 18 (S) | 26 (S) |
| Enteritidis 65/S/10 | 28 (S) | 28 (S) | 20 (S) | 26 (S) |
| Enteritidis 012PP2014 | 30 (S) | 30 (S) | 20 (S) | 28 (S) |

## Supplementary file 3

**The extended results of the bioinformatic analysis**

**Fig S3.**
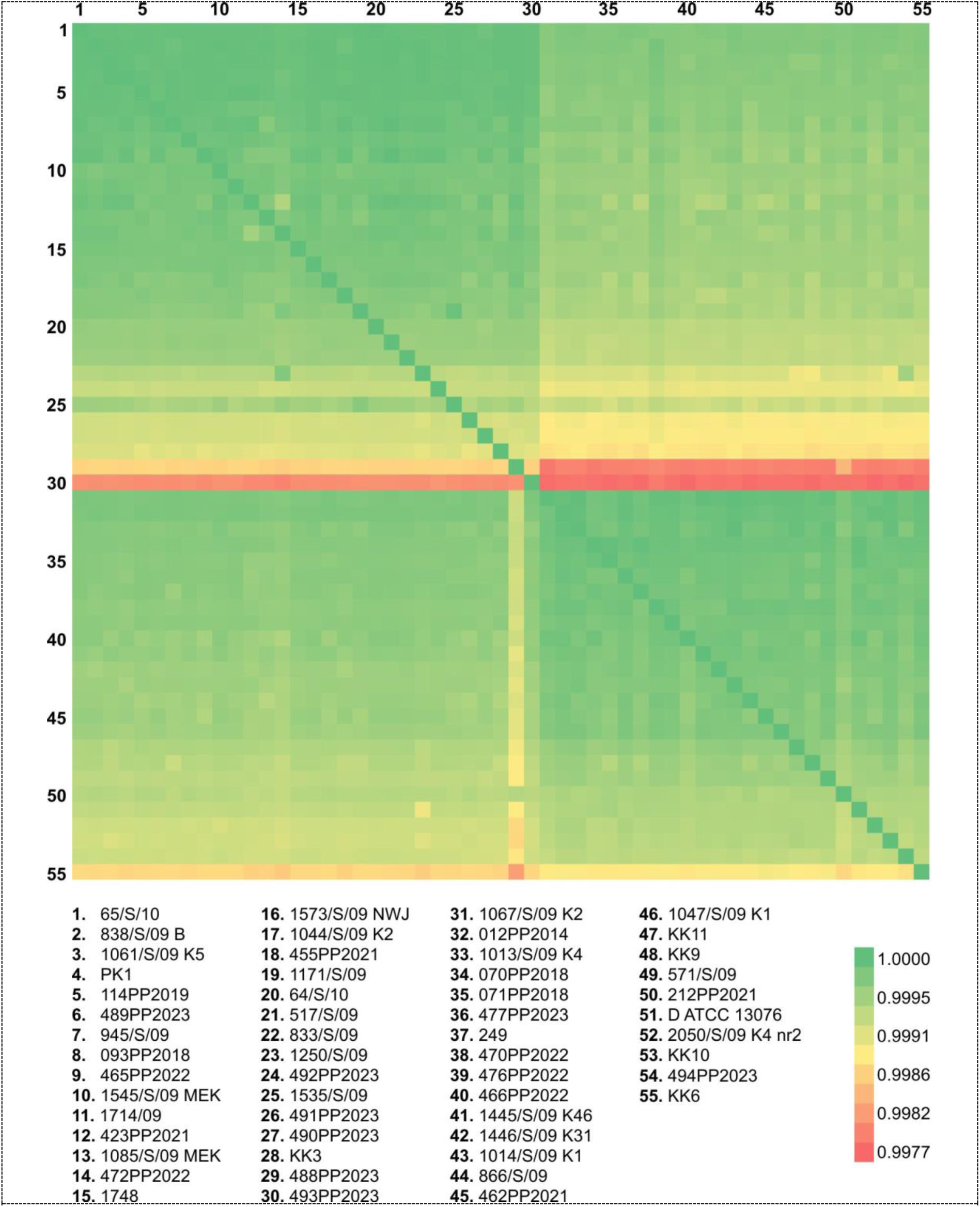
**ANI heatmap of the Enteritidis strains**

**Table S3.1.**
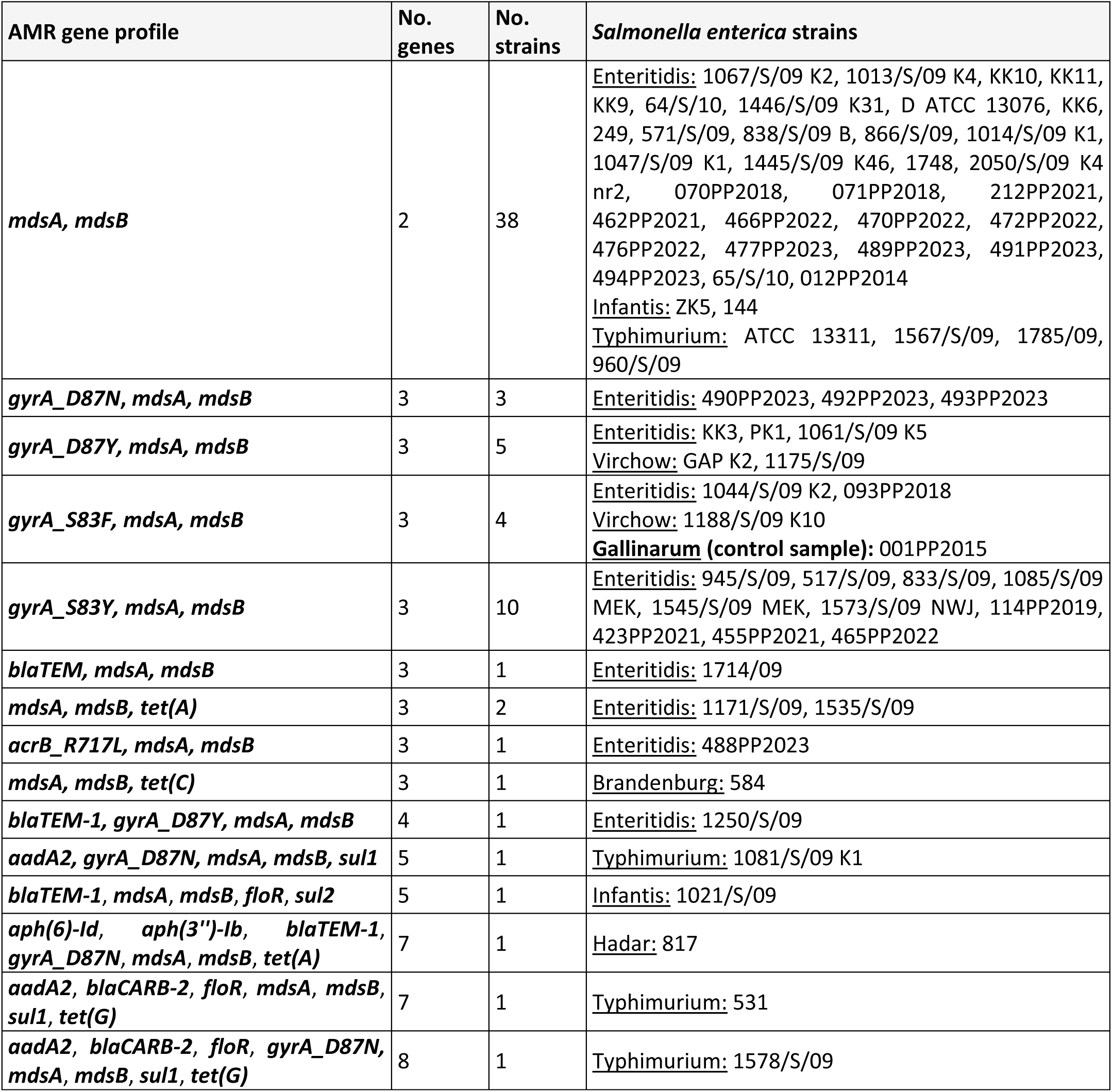
Putative antibiotic resistance genotypes in tested *Salmonella enterica* strains according to AMRFinderPlus analysis

For 71 out of 72 *Salmonella enterica* strains two multidrug efflux pump genes were identified - ***mdsA*** and ***mdsB***. These two genes were the only antibiotic resistance genes identified for the 32 *Salmonella enterica* ser. Enteritidis strains, 2 *Salmonella enterica* ser. Infantis and 4 *Salmonella enterica* ser. Typhimurium strains. On the other hand, there was one strain, which was found to be deprived of AMR genes: *Salmonella enterica* ser. Paratyphi A ATCC 19150.

The remaining tested *Salmonella enterica* strains revealed the presence of 3 to 8 AMR genes, amongst other: β-lactam antibiotic resistance genes - ***blaCARB-2, blaTEM, blaTEM-1*,** conferring resistance to streptomycin ***aph(6)-Id, aph(3’’)-Ib, aadA2*** genes, tetracycline resistance ***tet(A), tet(C), tet(G)*** genes, chloramphenicol/phenicol resistance ***floR*** gene, as well as ***sul1, sul2*** genes which confer resistance to sulfonamide antibiotics. There were also four different point mutations found in DNA gyrase subunit A **(*gyrA)*** gene, which correspond to quinolone (and/or triclosan) resistance, as well as unique (present for only one Egyptian *Salmonella enterica* ser. Enteritidis 488PP2023 strain (sample code: Enteritidis 488PP2023)) point mutation within multidrug efflux RND transporter permease subunit ***acrB*** gene conferring resistance to macrolide antibiotic azithromycin.

The highest number of 8 AMR genes was identified for *Salmonella enterica* ser. Typhimurium 1578/S/09 strain, for which analysis revealed presence of above mentioned ***mdsA*** and ***mdsB*** genes, **aadA2** gene (ANT(3’’)-Ia family aminoglycoside nucleotidyltransferase, conferring resistance to streptomycin), ***blaCARB-2*** gene (PSE family carbenicillin-hydrolyzing class A beta-lactamase, conferring resistance to β-lactam antibiotics), ***floR*** gene (chloramphenicol/florfenicol efflux MFS transporter), ***sul1*** (sulfonamide-resistant dihydropteroate synthase) and ***tet(G)*** gene (tetracycline efflux MFS transporter).

### Virulence profiles

Virulence factors profiles that were identified by VFDB for individual strains, were subsequently analysed within serovars.

### Salmonella enterica ser. Enteritidis

For fifty-five *Salmonella enterica* ser. Enteritidis strains from 165 to 325 VF per strain were identified, with the median of 174 virulence factors per strain (**Table 6**., column VF; **Table 7**.). Majority of the strains of this serovar (n= 24; **Table S3.2.**) had **175** virulence factors identified, which are listed in **Table S3.3.**

**Table S3.2.** Number of virulence factors identified for 55 *Salmonella enterica* ser. Enteritidis strains analysed in the study. Underlined strain – see description below

| Sample code | No. strains | No. VF |
| --- | --- | --- |
| 494PP2023 | 1 | 165 |
| 472PP2022 | 1 | 166 |
| 1446/S/09 K31, 838/S/09 B, 2050/S/09 K4 nr2 | 3 | 173 |
| 1067/S/09 K2, 1013/S/09 K4, KK10, KK11, KK9, D ATCC 13076, KK6, 249, 571/S/09, 866/S/09, 1014/S/09 K1, 1047/S/09 K1, 1445/S/09 K46, 070PP2018, 071PP2018, 462PP2021, 466PP2022, 470PP2022, 477PP2023, 488PP2023, 489PP2023, 492PP2023, 012PP2014 | 23 | 174 |
| 1044/S/09 K2, 945/S/09, 1714/09, 517/S/09, KK3, PK1, 64/S/10, 833/S/09, 1061/S/09 K5, 1085/S/09 MEK, 1250/S/09, 1545/S/09 MEK, 1573/S/09 NWJ, 1748, 093PP2018, 114PP2019, <u>212PP2021</u>, 423PP2021, 455PP2021, 465PP2022, 490PP2023, 491PP2023, 493PP2023, 65/S/10 | 24 | 175 |
| 1171/S/09, 1535/S/09 | 2 | 178 |
| 476PP2022 | 1 | 325 |

**Table S3.3.** 175 virulence factor related genes, along with virulence class identified for the majority (n=24, for sample codes see Table 6.) of *Salmonella enterica* ser. Enteritidis strains

| VFclass | Virulence factors | Related genes |
| --- | --- | --- |
| Fimbrial adherence determinants | Agf/Csg | *csgA, csgB, csgC, csgD, csgE, csgF, csgG* |
|  | Bcf | *bcfA, bcfB, bcfC, bcfD, bcfE, bcfF, bcfG* |
|  | Fim | *fimA, fimC, fimD, fimF, fimH, fimI, fimW, fimY, fimZ* |
|  | Lpf | *lpfA, lpfB, lpfC, lpfD, lpfE* |
|  | Pef | *pefA, pefB, pefC, pefD* |
|  | Peg | *pegA, pegB, pegC, pegD* |
|  | Saf | *safA, safB, safC, safD* |
|  | Sef | *sefA, sefB, sefC, sefD* |
|  | Stb | *stbA, stbB, stbC, stbD, stbE* |
|  | Std | *stdA, stdB, stdC* |
|  | Ste | *steA, steB, steC, steD, steE, steF* |
|  | Stf | *stfA, stfC, stfD, stfE, stfF, stfG* |
|  | Sth | *sthA, sthB, sthC, sthD, sthE* |
|  | Sti | *stiA, stiB, stiC, stiH* |
| Macrophage inducible genes | Mig-14 | *mig-14* |
|  | Mig-5 | *mig-5* |
| Magnesium uptake | Mg<sup>2+</sup> transport | *mgtB, mgtC* |
| Nonfimbrial adherence determinants | MisL | *misL* |
|  | RatB | *ratB* |
|  | ShdA | *shdA* |
|  | SinH | *sinH* |
| Regulation | PhoPQ | *phoP, phoQ* |
| Secretion system | TTSS (SPI-1 encode) | *hilA, hilC, hilD, iacP, iagB, invA, invB, invC, invE, invF, invG, invH, invI, invJ, orgA, orgB, orgC, prgH, prgI, prgJ, prgK, sicA, sicP, sipD, spaO, spaP, spaQ, spaR, spaS, sprB* |
|  | TTSS (SPI-2 encode) | *ssaC, ssaD, ssaE, ssaG, ssaH, ssal, ssaJ, ssaK, ssaL, ssaM, ssaN, ssaO, ssaP, ssaQ, ssaR, ssaS, ssaT, ssaU, ssaV, sscA, sscB, sseA, sseB, sseC, sseD, sseE, ssrA, ssrB* |
|  | TTSS effectors translocated via both systems | *slrP* |
|  | TTSS-1 translocated effectors | *avrA, sipA, sipB, sipC, sopA, sopB/sigD, sopD, sopE2, sopE, sptP* |
|  | TTSS-2 translocated effectors | *pipB2, pipB, sifA, sifB, spiC/ssaB, spvC, spvD, sseF, sseG, sseI/srfH, sseJ, sseK1, sseK2, sseL, sspH2* |
|  | ACE T6SS (Escherichia) | *aec31* |
|  |  | *aec32* |
| Serum resistance | Rck | *rck* |
| Stress adaptation | SodCI | *sodCI* |
| Toxin | SpvB | *spvB* |
| Adherence | LPS O-antigen<br>(*P. aeruginosa*) | - |
| Autotransporter | EhaB | *ehaB* |
| Immune evasion | LPS glucosylation<br>(*Shigella*) | *gtrA* |

The only exception in the above given VF contents turned out to be strain *Salmonella enterica* ser. Enteritidis 212PP2021 (denoted with underline in **Table 6**.), for which different TTSS-2 translocated effector coding gene was identified (***gogB*** gene instead of *sseK2*). *GogB* is reported to be an anti-inflammatory effector that helps regulate inflammation-enhanced colonization by limiting tissue damage during infection [Pilar et al. GogB is an anti-inflammatory effector that limits tissue damage during *Salmonella* infection through interaction with human FBXO22 and Skp1. PLoS Pathog. 2012;8(6):e1002773]

Out of the group of 23 strains, for which 174 VF were identified, 20 strains were deprived of *sseK2* gene, 1 strain (Enteritidis 489PP2023) of fimbrial adherence ***sefD*** gene, 1 strain (Enteritidis 492PP2023) of secretion system ***hilA*** gene and 1 strain (Enteritidis 488PP2023) of ***gtrA*** gene associated with immune evasion in *Shigella*.

As for the strains for which 178 virulence factors were identified (Enteritidis 1171/S/09, Enteritidis 1535/S/09), the additional genes were associated with adherence type IV pili typical for *Yersinia* - ***pilQ, pilR, pilS, pilW.*** On the other hand these two strains were deprived of specific to *Escherichia* ACE T6SS - ***aec32*** gene.

The strain Enteritidis 472PP2022, for which analysis revealed the presence of only 166 virulence associated genes, turned out to lack: fimbrial adherence pef genes (***pefA, pefB, pefC, pefD***), macrophage inducible gene ***mig-5***, and TTSS-2 translocated effector genes (***spvC, spvD***). Strain Enteritidis 494PP2023 was additionally deficient of *sseK2* gene.

In contrast, strain Enteritidis 476PP2022, for which 325 virulence factors were identified was checked for possible contamination but none was determined (over 98% of sequencing reads assigned to *Salmonella* genus, no 16S RNA fragments different from *Salmonella*). The additional virulence factors genes were associated with regulation, secretion systems, serum resistance, stress adaptation, toxin, adherence, anaerobic respiration, antiphagocytosis, autotransporter, biofilm formation, cell surface components, copper uptake, efflux pump, endotoxin, enzyme, immune evasion, invasion, iron uptake, lipid and fatty acid metabolism, motility, nutritional factor, nutritional virulence, protease, quorum sensing, serum resistance and immune evasion.

### Salmonella enterica ser. Infantis

For *Salmonella enterica* ser. Infantis analysis revealed 151 virulence factors for strains denoted Infantis ZK5, Infantis 144 and 155 genes for strain Infantis 1021/S/09.

When compared with serovar Enteritidis, these strains are lacking genes encoding Pef, Peg Sef, and Ste fimbrial adherence determinants, macrophage inducible gene *mig5*, serum resistance gene *Rck,* stress adaptation gene *SodCI* and *spvB* toxin gene. Infantis on the other hand has Stc and Tcf fimbrial adherence determinants and O-antigen (*Yersinia*), when compared with Enteritidis serovar.

The higher number of virulence factors that were identified for strain *Salmonella enterica* ser. Infantis 1021/S/09, is due to the adherence type IV pili (*Yersinia*) genes: ***pilQ, pilR, pilS, pilV, pilW***, however it is lacking secretion system ***ssrA*** gene.

### Salmonella enterica ser. Typhimurium

Seven *Salmonella enterica* ser. Typhimurium strains in the analysed dataset, were found to have variable number of virulence factors. There were 153 such genes identified for strain Typhimurium ATCC 13311, 165 for strains Typhimurium 1567/S/09, Typhimurium 1785/09 and Typhimurium 960/S/09, 167 genes for strains Typhimurium 1578/S/09 and Typhimurium 531, and the highest number of 168 genes for strain Typhimurium 1081/S/09 K1. Genes diversifying the strains are given in **Table S3.4.**

**Table S3.4.** Virulence factors diversifying *Salmonella enterica* ser. Typhimurium strains

| VFclass | Virulence factors | Related genes | 1081/S/09 K1 | 1578/S/09 | 531 | 1567/S/09 | 1785/09 | 960/S/09 | ATCC 13311 |
| --- | --- | --- | --- | --- | --- | --- | --- | --- | --- |
|  |  |  | 168 | 167 | 167 | 165 | 165 | 165 | 153 |
| Fimbrial adherence determinants | Pef | pefA | + | + | + | + | + | + | - |
|  |  | pefB | + | + | + | + | + | + | - |
|  |  | pefC | + | + | + | + | + | + | - |
|  |  | pefD | + | + | + | + | + | + | - |
|  | Saf | safA | + | + | + | + | + | + | - |
|  |  | safB | + | + | + | + | + | + | - |
|  |  | safC | + | + | + | + | + | + | - |
|  | Stf | stfE | + | + | + | + | - | - | + |
| Macrophage inducible genes | Mig-5 | mig-5 | + | + | + | + | + | + | - |
| Nonfimbrial adherence determinants | ShdA | shdA | + | + | + | + | + | + | - |
| Secretion system | TTSS-2 translocated effectors | spvC | + | + | + | + | + | + | - |
|  |  | spvD | + | + | + | + | + | + | - |
|  | Lvh (Legionella vir homologs) type IVA secretion system | traG | + | - | - | - | - | - | - |
| Serum resistance | Rck | rck | + | + | + | + | + | + | - |
| Toxin | SpvB | spvB | + | + | + | + | + | + | - |
|  | Typhoid toxin | pltA | + | + | + | - | - | - | + |
| Immune evasion | LPS glucosylation (Shigella) | gtrA | + | + | + | - | + | + | - |

In comparison with Enteritidis strains, Typhimurium strains in the collection are deprived of Peg, Sef, Ste fimbrial adherence determinants genes and secretion system gene *sopE.* Conversely to Enteritidis serovar, all Typhimurium strains have Stc and Stj fimbrial adherence determinants genes as well as secretion system gene *sseK2*.

### Salmonella enterica ser. Virchow

All three *Salmonella enterica* ser. Virchow strains have similar number of virulence factors (from 169 to 170) and differ only in terms of the presence of O-antigen (*Yersinia*) in strains 46/182/23 and 47/182/23.

When compared with Enteritidis serovar strains, Virchow representatives in the study are deprived of Pef, Peg and Sef fimbrial adherence determinants genes, macrophage inducible gene *mig5*, nonfimbrial adherence gene *ratB,* serum resistance gene *Rck,* stress adaptation gene *SodCI* and *spvB* toxin gene.

On the other hand, Virchow strains have Stc, Stj, Stk and Tcf fimbrial adherence determinants and O-antigen (*Yersinia*), when compared with Enteritidis serovar.

### Salmonella enterica ser. Hadar

For the only representative of Hadar serovar (Hadar 817) 166 virulence factor genes were identified. As compared with Enteritidis strains, it was found to lack genes encoding Pef, Peg and Sef fimbrial adherence determinants, macrophage inducible gene *mig5*, nonfimbrial adherence gene *ratB,* serum resistance gene *Rck,* stress adaptation gene *SodCI* and *spvB* toxin gene.

Opposite to Enteritidis, for Hadar strain Stc, Stj and Stk fimbrial adherence determinants and O-antigen (*Yersinia*), were identified.

### Salmonella enterica ser. Branderburg

*Salmonella enterica* ser. Brandenburg representative (Brandenburg 584) was found to have 165 virulence factor genes. Comparison with Enteritidis strains, revealed only one of Lpf and Pef fimbrial adherence determinants genes (*lpfC* and *pefB*, respectively), whereas Sef, Ste and Stf genes were absent at all, just as macrophage inducible gene *mig5*, nonfimbrial adherence gene *ratB*, serum resistance gene *rck*, stress adaptation gene *SodCI, spvB* toxin gene and secretion system genes *sopE, spvC, spvD* and *sseI/srfH*. Contrary to Enteritidis strains present were all 7 Sta, 5 Stj and 4 Tcf fimbrial adherence determinants genes, three typhoid toxin genes *cdtB, pltA,* and *pltB*, as well as seven *fae* genes associated with K88 fimbriae in *Escherichia*.

### Salmonella enterica ser. Paratyphi A

For the only *Salmonella enterica* ser. Paratyphi A (Paratyphi A ATCC 19150) in the collection 160 VF were identified. In comparison with Enteritidis it was lacking genes encoding Pef, Peg and Sti fimbrial adherence determinants, macrophage inducible gene *mig5,* serum resistance gene *Rck,* stress adaptation gene *SodCI, spvB* toxin gene and secretion system genes *avrA, spvC, spvD, sseI/srfH, sseJ, sseK1, sspH2.* As compared with Enteritidis present were Stk and Tcf fimbrial adherence determinants, three typhoid toxin genes (*cdtB, pltA, pltB*), as well as invasin A from *Yersinia*.

### *Salmonella enterica* ser. Gallinarum control strain

*Salmonella enterica* ser. Gallinarum control strain (Gallinarum 001PP2015) was found to have 172 virulence factor genes. When compared with Enteritidis, it turns out that no Pef and Std, fimbrial adherence determinants, macrophage inducible gene *mig5,* nonfimbrial adherence gene *ratB,* serum resistance gene *rck*, *spvB* toxin gene and secretion system genes *spvC, spvD, sseI/srfH, sspH2* were identified. On the other hand, for the strain 17 ACE T6SS (*Escherichia*) genes were identified (***aec15, aec16, aec17, aec18, aec19, aec22, aec23, aec24, aec25, aec26, aec27/clpV, aec28, aec29, aec30, aec31, aec32***), whereas for Enteritidis strains maximum three of them were identified (***aec30, aec31, aec32***).

